# 4polar-STORM polarized super-resolution imaging of actin filament organization in cells

**DOI:** 10.1101/2021.03.17.435879

**Authors:** Caio Vaz Rimoli, Cesar Augusto Valades Cruz, Valentina Curcio, Manos Mavrakis, Sophie Brasselet

## Abstract

Advances in single-molecule localization microscopy are providing unprecedented insights into the nanometer-scale organization of protein assemblies in cells and thus a powerful means for interrogating biological function. However, localization imaging alone does not contain information on protein conformation and orientation, which constitute additional key signatures of protein function. Here, we present a new microscopy method which combines for the first time Stochastic Optical Reconstruction Microscopy (STORM) super-resolution imaging with single molecule orientation and wobbling measurements using a four polarization-resolved image splitting scheme. This new method, called 4polar-STORM, allows us to determine both single molecule localization and orientation in 2D and to infer their 3D orientation, and is compatible with high labelling densities and thus ideally placed for the determination of the organization of dense protein assemblies in cells. We demonstrate the potential of this new method by studying the nanometer-scale organization of dense actin filament assemblies driving cell adhesion and motility, and reveal bimodal distributions of actin filament orientations in the lamellipodium, which were previously only observed in electron microscopy studies. 4polar-STORM is fully compatible with 3D localization schemes and amenable to live-cell observations, and thus promises to provide new functional readouts by enabling nanometer-scale studies of orientational dynamics in a cellular context.

## Introduction

Protein conformation and the precise way in which proteins arrange in space to form higher-order macromolecular assemblies are key elements of biological functions in cells and tissues. Adhesion of animal cells to the extracellular matrix is driven, for example, by dramatic conformational changes in force-sensing and force-transducing proteins such as integrins and talins. The precise geometry of actin filament assemblies^1,2,3,4,5^ and its remodelling in space and time are further determinant for cell mechanics driving essential biological processes, including immune responses and tissue development. Thus, understanding the function of a protein and its interaction with its partners, necessitates that we observe its organization at the nanometer scale, both in position and orientation. This need is shared by many fields in biology from immunology, neurobiology and mechanobiology, to developmental biology. Current methods reporting protein organization such as electron microscopy or X-ray diffraction are however not yet applicable to a live imaging context. Single molecule localization microscopy (SMLM) has brought considerable progress towards this goal, enabling imaging with a resolution down to tens of nanometers even in live cells^6,7,8,9^. However, while these methods report the localization of single molecules with high precision, they do not measure their orientation. If fluorophores are linked to the proteins of interest rigidly enough^10^, reporting their orientation could provide precious information on the structural organization of the attached proteins and on their conformational behaviors, which is inherently missing in localization-based optical imaging methods. While measuring fluorophore orientation precisely and accurately is a challenge that is of high interest in the field of SMLM imaging, it is however a delicate task. First, it is necessary to not only measure their 3D orientation, averaged over the imaging time, but also the extent of their orientation fluctuations, which naturally occurs when fluorophores wobble at fast time scales^10^ (Fig. 1a). A failure to uncouple their mean orientation from their fluctuations makes it impossible to determine accurately how the fluorophore-conjugated molecules are organized and can lead to misleading interpretations^10^. Second, orientation and spatial position are difficult to disentangle in SMLM, because of their intrinsic coupling in the process of the formation of their PSF image in a microscope^11,12^. The development of an optimal method to disentangle spatial position, mean orientation and orientation fluctuations is still an ongoing research^13^. One approach is to encode the orientation and wobbling information into the shape of the single molecules’ point spread function (PSF), using custom-designed phase or birefringent masks^14,15,16^. This strategy comes however at the price of some constraints, which restrain its use in regular SMLM imaging. PSF engineering induces an increase of the PSF size, with the PSF shape including both orientation and wobbling information in an intricate way, limiting its use in densely-labelled structures. PSF engineering also involves practical and methodology difficulties due to complex data analysis procedures and stringent calibrations to avoid sources of bias such as optical aberrations. Along similar lines, exploiting un-engineered PSF shape changes due to image defocusing^17,18^ or to the proximity of the molecule to the coverslip interface^19^ has been proposed, however with similar limitations as encountered in PSF engineering. A less constraining approach is to use polarization projections of the image plane, and perform ratiometric intensity measurements between different polarization channels. Two-orthogonal polarization splitting has allowed fluorescence anisotropy measurements in isotropic environments^20^ and used to quantify actin filament alignment in 2D^10^, however with the inconvenience of an estimation ambiguity for fluorophore orientations symmetric relative to the polarization axes. Additionally, two-orthogonal polarization splitting does not efficiently decouple orientation from wobbling. Both ambiguities have been waived by the use of a four-polarization projection scheme^21,22,23,24^. A strong limitation still present in currently reported four-polarization split approaches is however that due to the high numerical apertures used, the intensities measured are strongly influenced by the off-plane 3D orientation of the fluorophores, resulting in large inaccuracies in the determination of their wobbling^23^. The failure to provide accurate measurements of fluorophore wobbling has precluded its use as an additional readout for protein organization. It is indeed conceivable that fluorophore wobbling is related to local packing constraints of the labelled protein. Importantly, single molecule studies using four polarizations projections have not yet been applied to super-resolution imaging and have so far been limited to situations employing sparse labelling or/and photobleaching to obtain single fluorophores^21,22,23,24^. In this work, we combine for the first time Stochastic Optical Reconstruction Microscopy (STORM) super-resolution imaging with single molecule orientation and wobbling measurements using four-polarization image splitting in a new method called 4polar-STORM. 4polar-STORM imaging is compatible with high labelling densities and is thus ideally placed for the determination of the organization of dense protein assemblies in cells. 4polar-STORM imaging further uses a slightly reduced detection numerical aperture, allowing us to not only determine the 2D mean orientation and wobbling of fluorophores in a reliable way, but also to infer their 3D orientation and thus provide additional information on protein organization and function that is otherwise difficult to obtain.

**Figure 1.**
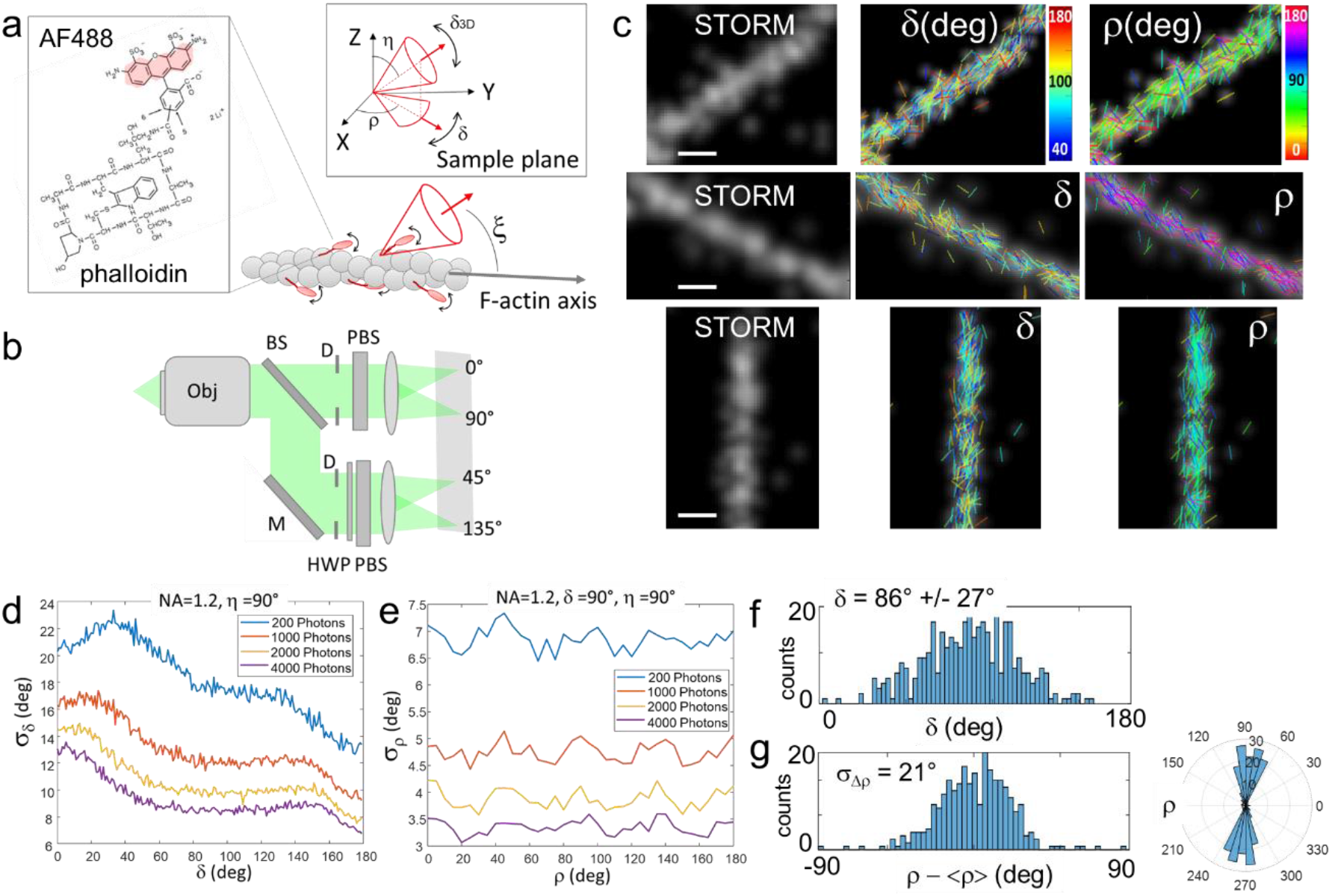
(a) Schematic representation of a wobbling Alexa Fluor 488 (AF488)-phalloidin conjugate labelling an actin filament (F-actin). The structure of the phalloidin conjugate is shown on the left, with the fluorophore moiety highlighted in red. (ρ,η) represent the mean orientation of a single fluorophore in 3D, with ρ the projected orientation in the sample plane (XY) and η its off-plane tilt angle with respect to the optical axis of the microscope (Z). δ_3D_ is the wobbling cone angle of the fluorophore in 3D, and δ its projection in the sample plane. ξ is the mean orientation of the fluorophore relative to the actin filament axis. (b) Schematic optical setup of 4polar-STORM imaging. BS, beam splitter; M, mirror; D, diaphragm; HWP, half-wave plate; PBS, polarizing beam splitter. (c) Examples of 4polar-STORM images of single AF488-phalloidin molecules labelling single actin filaments. Left panels depict single molecule localization STORM images (blurred using a Gaussian filter with a sigma of 0.3 pixel size). Middle and right panels depict single molecule wobbling (δ) and orientation (ρ) measurements overlaid with the STORM image depicted in grayscale. Each single molecule is represented as a stick whose orientation is ρ relative to the horizontal axis and whose color is the measured parameter (ρ or δ). Scale bars, 170 nm. (d) Monte Carlo simulations of the expected precision on δ under different signal level conditions (signal in photons, 1000 photons = 4.5×10^4^ camera counts). 5000 realizations are used for η = 90° (in-plane molecules) in the presence of camera detection noise using the experimental conditions and a gain of 300 (the results do not depend on ρ) (see Supplementary Fig. S3). (e) Corresponding Monte Carlo simulations for the estimation of ρ precisions reached for (δ = 90°, η = 90°) (the results do not strongly depend on δ apart for δ = 0° and 180°). (f) Representative experimental δ histogram obtained on a straight segment of a single actin filament (see (c)). (g) Corresponding ρ histogram in both standard and polar-plot representations, depicting the distribution of orientation angle values relative to the average within the measured region of interest, Δρ = ρ –<ρ>. σ_Δρ_ is the standard deviation of Δρ. For both histograms, intensities are thresholded above 5×10^4^ camera counts and localization precisions are thresholded below 0.15 pixels (σ_loc_ < 20 nm).

With this new approach, we reveal the nanometric-scale structural organization of actin filaments inside dense actin filament-based structures involved in the adhesion and motility of cells, notably stress fibers (SFs) and lamellipodia. Actin is at the center of interest in the development of super-resolution imaging methods where so far, only localization-based images have been exploited^25,26,27,28,29,30^. Here we use 4polar-STORM imaging to quantify the orientational behaviour of fluorophores in single actin filaments used as a reference, and in fixed cells. We show that actin filaments are highly aligned in all types of SFs in cells, in line with EM studies^31,32^, and further reveal that SFs contain not only 2D but also 3D oriented populations of actin filaments, whose nanometer-scale organization is consistent with the different proposed mechanisms for their assembly. Mild pharmacological inhibition of myosin II activity to relax contractile SFs, while keeping SFs macroscopically intact, led to a perturbation of the nanometric actin filament organization, as revealed by 4polar-STORM imaging, highlighting the dependence of contractile SF organization on Myosin II activity. Finally, 4polar-STORM imaging in the dense meshwork of the lamellipodium at the leading edge of motile cells revealed bimodal distributions of actin filament orientations, with non-negligible 3D oriented filament populations. Such bimodal orientation distributions have not been reported with other super-resolution light microscopy methods, to our knowledge, due to the high actin density in this area^25^, and were previously only observed in EM studies of the lamellipodium^3,33,34,35,36^. This new method is amenable to live-cell observations, and promises to complement electron microscopy studies in cells, while allowing for nanometer-scale measurements of molecular organization in large (tens of micrometers) fields of views.

### 4polar-STORM imaging applied to actin filaments

Fluorophores attached to a protein act as emission dipoles. While their position is directly defined as the center of their PSF image, their orientation is not directly extractable. A fluorophore is represented by its mean orientation (ρ,η) averaged over the imaging integration time, and its wobbling angle (δ_3D_) explored during this integration time (Fig. 1a). 4polar-STORM measures the fluorophore’s orientation and wobbling projected in the sample plane (ρ,δ) (Fig. 1a) based on the projection of the fluorescence signal on four polarizations channels along the directions 0°, 45°, 90°, and 135° respectively (0° corresponding here to the horizontal direction of the sample) (Fig. 1b). The 4-polarization splitting approach leads to only a minor deformation and enlargement of the image PSF, affecting minimally the signal to noise ratio (SNR) and PSF complexity. In contrast to a 2-polarization projection^10^, 4-polarization projection permits to waive coupling ambiguities and therefore to retrieve ρ and δ independently. Supplementary Note 1 details the theoretical framework for the retrieval of the orientation parameters (ρ,δ). We note that δ is the sample-plane 2D projection of the wobbling cone angle δ_3D_, it therefore differs from the real 3D wobbling value δ_3D_. At large off-plane tilt angles in particular (small η angle in Fig. 1a), the projection of the wobbling cone angle is biased and δ is an overestimation of δ_3D_. Theoretical calculations of the dependence of this bias on the detection numerical aperture (NA) and on the tilt angle of the fluorophore, show that a solution for minimizing this bias is to lower the NA to a value close to 1.2 (Supplementary Note 1). Even though both SNR and PSF size are expected to be slightly degraded for this lower NA, this permits to give a low-bias estimate of δ_3D_ with a reasonable compromise on the loss of signal, as long as the tilt angle of the fluorophore off-plane orientation η does not surpass 45° (Supplementary Note 1 and Fig. S1).

The retrieval of the orientation parameters (ρ,δ) consists in detecting first the 2D position of single molecules in each of the polarization channels, previously corrected for imperfections in the detection paths (Supplementary Note 2) and registered (Supplementary Note 3). Each molecule is associated to its pairs in all polarization channels, where polarized PSF amplitudes and sizes are deduced from a Gaussian fit in order to calculate intensities along each of the polarized channels (Fig. 1b) (Supplementary Note 3). We note that if molecules have a well-defined orientation and are oriented off-plane, their PSF will enlarge and deform towards a donut-shape^37^. Even though this effect is minimized when molecules wobble^38^, the PSF fit has to allow for larger ranges of sizes, since the measured PSF size for tilted molecules can be larger than for in-plane molecules. Once intensities are determined in the four polarization channels, a retrieval calculation permits to extract both ρ and δ parameters per molecule (Supplementary Note 1). This calculation uses a relatively simple model, for computational speed reasons, which is shown to be very close to a complete model characterization accounting for the inversion of the propagation equations (Supplementary Note 1 and Fig. S2).

Using the theoretical framework we established for the retrieval of orientation parameters, we aimed at using 4polar-STORM imaging to measure the nanometer-scale actin filament organization in complex assemblies in cells labelled with fluorophore-conjugated phalloidin molecules, which provide specific labelling of actin filaments^39,40^. Our measurements will allow us to extract both the degree of angular fluctuations of the fluorophores (δ angle in Fig. 1a), and their mean orientation in the sample plane (ρ angle in Fig. 1a). To evaluate the fluorophore orientation behaviour in a flat, single actin filament with a well-defined direction, we started by reconstituting single actin filaments immobilized on a glass surface (see Methods). This provides a reference for later deciphering actin filament organization in unknown, more complex assemblies. Figure 1c shows the results obtained on isolated actin filaments labelled with Alexa Fluor 488 (AF488)-phalloidin. Single AF488 molecules measured by 4polar-STORM are represented as sticks whose color is the wobbling angle δ and whose orientation, relative to the horizontal axis, is the angle ρ. Figure 1c shows that AF488 molecules are oriented along the actin filament axis, and exhibit a non-negligible degree of orientational flexibility δ. The expected precision on both ρ and δ, depicted in Fig. 1d,e, was evaluated by Monte Carlo simulations under different signal levels, accounting for the camera noise (Supplementary Figure S3). Typical measured total intensities from single molecules are around 5 to 10×10^4^ camera counts. At 5×10^4^ camera counts (e.g. 1100 photons), we expect an error of 12° for δ in the measured range of δ values (Fig. 1d), and of 5° degrees for ρ (Fig. 1e). This error decreases with the signal level and is expected to increase in the presence of background. In order to exclude degraded signal in 4polar-STORM due to background conditions, and provide high precision estimates, we systematically threshold the experimental data to a minimum intensity of 5×10^4^ camera counts, and to a maximum localization precision value σ_loc_ of 0.15 pixels (corresponding to 20 nm). Straight segments of actin filaments were selected under such conditions, and typical statistics obtained on δ and ρ in these regions are plotted in Fig. 1f and Fig. 1g respectively. Measurements of the orientation angles ρ are depicted relatively to the average <ρ> over the molecules measured in the region of interest, Δρ = ρ − <ρ> (Fig. 1g). The distribution of Δρ is characterized by a standard deviation σ_Δρ_ which represents the range of orientations explored by single AF488 molecules with respect to the actin filament; σ_Δρ_ also gives a measurement of the fluorophore tilt angle with respect to the filament axis (*ξ* angle in Fig. 1a). Figures 1f,g show that AF488 labels are, on average, oriented along the actin filament direction with a tilt angle of about *ξ* = 20° with respect to the actin filament axis, and a non-negligible wobbling of δ = 85-90°. This is consistent with the fact that the phalloidin-fluorophore conjugate exhibits a very small size (on the order of 1 nm) and a structure that fits in the groove formed by three neighbouring G-actin monomers^41^, while leaving some space for mobility for the fluorophore. As previous studies have suggested, this degree of mobility is expected to originate mostly from the structure of the fluorophore itself and its precise conjugation to the phalloidin moiety, AF488 being among the least wobbly fluorophores among STORM dyes^10^.

Using the same labelling approach, we next investigated the organization of actin filaments in structures that are expected to be highly organized in cells, notably actin stress fibers (SFs). We focused on ventral stress fibers, both ends of which associate with focal adhesions (FAs) on the ventral surface of the cell; on dorsal stress fibers, with one end associating with FAs on the ventral surface and the other end extending upwards toward the dorsal cell surface; and on meshworks (Fig. 2a). A minimum intensity threshold of 5×10^4^ camera counts and a maximum localization precision value σ_loc_ of 0.15 pixels are applied to all data in order to exclude estimates that lead to low precision and inaccuracy in the parameters’ determination. 4polar-STORM images of an actin-stained cell are shown in Fig. 2b-d, which depict respectively the single molecule localization image (STORM), the ρ and the δ images from the same cell. In well isolated thin ventral SFs, AF488 is oriented predominantly along the SF direction (ROI 1 in Figs. 2e,f, with polar histograms of single molecule orientations shown as insets in f), similarly to what was observed in single actin filaments (Fig. 1). This observation confirms that these SFs are made of highly parallel filaments, which is expected from the tight crosslinking of these structures^3^. Many molecules exhibit however larger δ values than the ones measured in single filaments. This is even more pronounced in ventral SF parts close to FAs (ROI 2 in Figs. 2e,f) or in dorsal SFs (ROI 3 in Figs. 2e,f); larger δ values appear as progressively redder sticks in the δ images. In these regions, ρ angles also distribute over a large range of orientations, not necessarily along the SF (see zooms in Fig. 2g, compare ROI 6 with ROI 5) and δ angle distributions shift to high values (see polar histogram insets in Figs. 2f and histograms in Fig. 2h, compare ROIs 2-3 with ROI 1). This trend was observed for multiple SFs in all measured cells (see other examples in Supplementary figure S4). At last, in seemingly homogeneous meshworks, AF488 orientations ρ seem to follow preferential directions with much larger distributions (ROI 4 in Figs. 2e,f,h).

**Figure 2.**
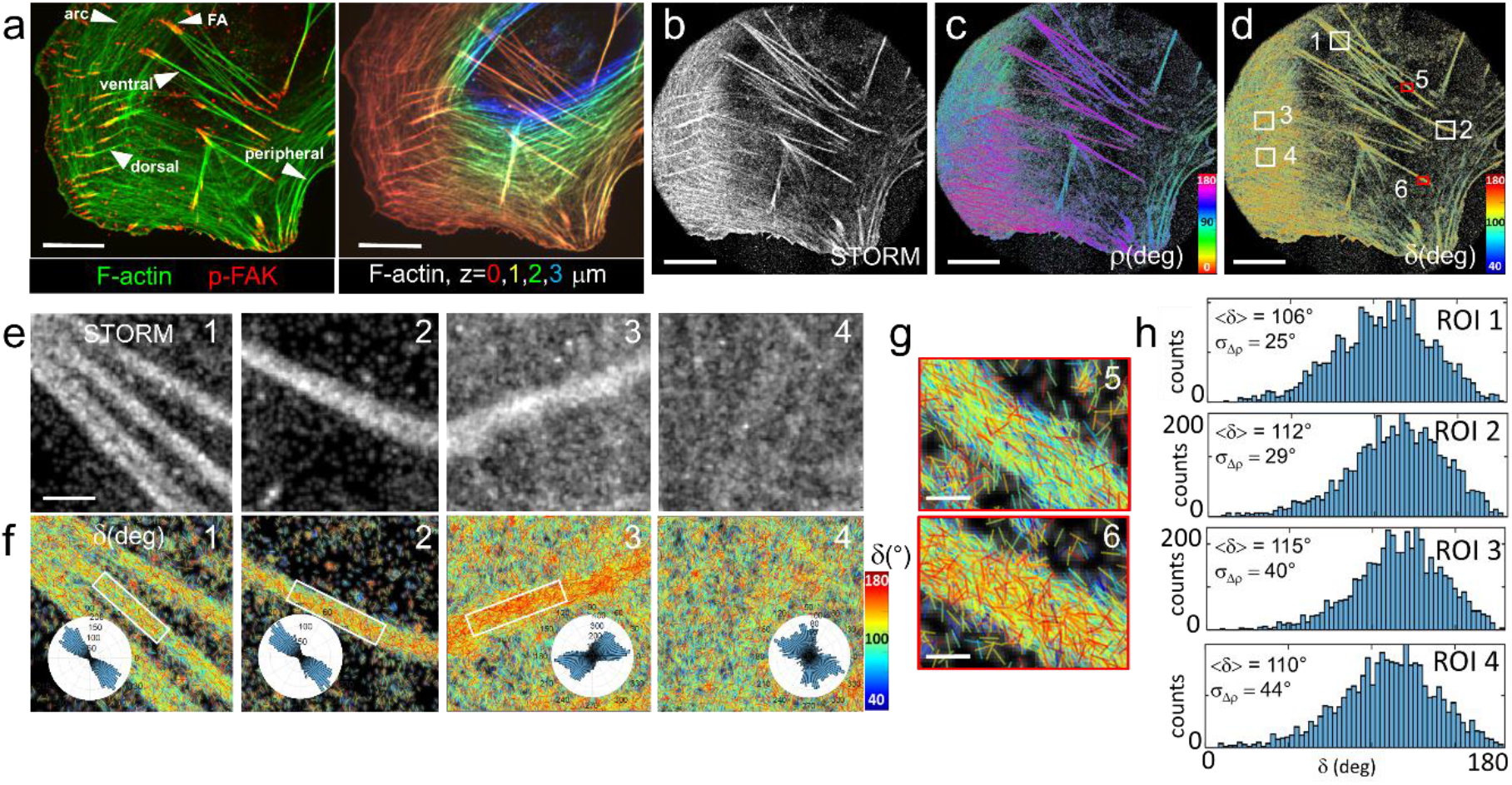
4polar-STORM imaging of actin filament organization in fixed U2OS cells. (a) Left: Spinning-disk fluorescence image of a U2OS cell (green, AF488-phalloidin-labelled F-actin; red, p-FAK). White arrows indicate focal adhesions (FAs) and the types of stress fibers (SFs) of interest. Right: z-stack mage of F-actin, z is color-coded as indicated. (b) Large field of view single molecule (AF488) localization STORM image of the same cell. (c) Corresponding 4polar-STORM ρ stick image with color-coded orientation measurements. (d) Corresponding 4polar-STORM δ stick image with color-coded wobbling angle measurements. (e) STORM and (f) corresponding δ stick images of zoomed regions of interest (ROI) (see squares in (d)). ROI 1, ventral SF; ROI 2, FA; ROI 3, dorsal SF; ROI 4, meshwork. Insets in (f) depict the polar-plot histograms of ρ in the rectangle region indicated (whole region for ROI 4). (g) δ stick images of zoomed ROIs 5 and 6 (see red squares in (d)) exhibiting respectively low δ and high δ populations. ROI 5, ventral SF; ROI 6, FA. (h) Histograms of δ values for ROIs 1-4. Values of <δ> (over all measured molecules) and of the standard deviation σ_Δρ_ of Δρ = ρ − < ρ > are shown. For all images, intensities are thresholded above 5×10^4^ camera counts and localization precisions are thresholded below 0.15 pixels (σ_loc_ < 20 nm). Scale bars (a-d), 7 μm; (e), 800 nm; (g), 260 nm.

To understand the presence of large δ values and wide ρ distributions in SFs, we investigated possible correlations between these two parameters. We found first that in all observed SFs, large δ values generally correlate with a wide distribution of ρ values (Fig. 3a). This behaviour could physically correspond to molecules that are freely, isotropically rotating, but we exclude such an effect. First, no free phalloidin-fluorophore conjugates are expected given the high affinity of phalloidin for actin filaments. Second, higher wobbling angles were not present in single actin filaments and thin actin bundles. We exclude also a sensitivity of wobbling to actin filament packing within bundles, considering the small size of phalloidin-AF488; single actin filaments do not present packing-related constraints and did not exhibit high wobbling angles. Our hypothesis for the observed high δ values is the presence of 3D oriented actin filaments (off-plane η angle in Fig. 1a) that lead to an overestimation of δ. The use of a relatively low detection NA of 1.2 minimizes the bias induced by 3D orientations as compared to higher NA conditions, but does not entirely exclude this effect. Typically, wobbling molecules with δ_3D_ = 90° oriented off-plane by η = 45° lead to a measured δ = 150° (Supplementary Fig. S1). 3D oriented filaments also naturally lead to larger deviations of ρ. Typically a ρ distribution of σ_Δρ_ = 20° for an in-plane filament would increase dramatically to an apparent σ_Δρ_ ~ 90° when this filament is tilted off-plane by 45° (Supplementary Fig. S5). To test the hypothesis of possible 3D oriented filaments, we investigated the correlation of δ and ρ with the single molecule detection parameters, in particular their intensity, which is expected to decrease when fluorophores are tilted off plane due to their lower photo-excitation, and the PSF radius, which is expected to increase when molecules are either tilted or out of focus. Large δ values and wider distributions of ρ values are indeed found to correlate with large PSF radii and with the lowest levels of intensities (Fig. 3b,c). Thresholding intensities and σ_loc_ values as above, as well as keeping PSF radii below 1.15 pixels (~150 nm) leads to a net reduction of the populations of high δ values (Fig. 3d; compare with Fig. 2f) (see also Supplementary Fig. S4). Larger populations of large PSF radii can also be found in SFs that are visibly enriched in high δ values (Supplementary Fig. S6). These populations are attributed to molecules tilted off plane (also possibly positioned at slightly different heights), and therefore to populations of filaments that are tilted with respect to the sample plane. Such tilted filaments are found, as expected, more frequently at the FA sites and in dorsal SFs, both of which are expected to contain off-plane filament populations, in contrast to ventral SFs, which lie in the plane of the substrate (Fig. 2a).

**Figure 3.**
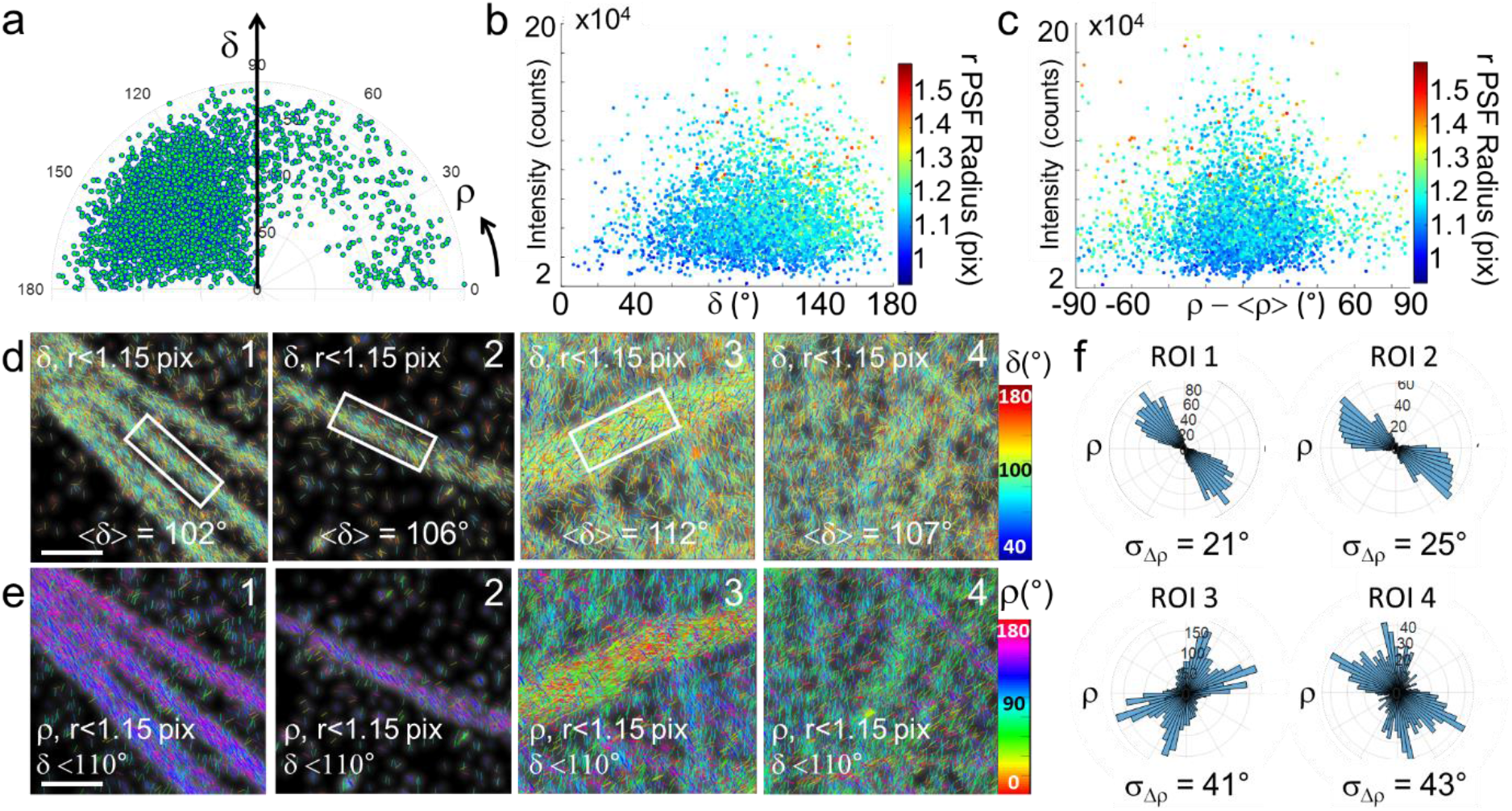
Influence of the detection parameters on 4polar-STORM imaging. (a) Scatter polar representation of (δ,ρ) values obtained in single AF488 molecules (all detected intensities) present in ROI 1 (rectangle in (d)). Each marker is a molecule present in the ROI, plotted using cylindrical coordinates δ as a radius and ρ as an angle respective to the horizontal axis. (b) Same δ values as in (a), using intensity counts as a vertical coordinate, and the PSF radius r as a color code for the markers. Each marker is a single molecule. (c) Same representation using, for the horizontal axis, ρ values centered with respect to their average orientation <ρ> (d) Same 4polar-STORM images as in Fig. 2e,f, depicting δ sticks only for molecules for which r < 1.15 pixels (149 nm). The corresponding average <δ> over all measured molecules (present in the rectangle for ROIs 1-3 or the whole region for ROI 4) are shown. (e) Same ROIs as in (d) showing ρ stick images only for molecules with δ < 110°. (f) Corresponding polar-plot histograms of ρ for regions indicated in (e). The corresponding σ_Δρ_ values are shown. Scale bar (d-e), 800 nm.

Estimation of the off-plane angle can be done qualitatively from the theoretical δ vs η bias dependence (Supplementary Fig. S1). In the case of actin filaments labelled with AF488-phalloidin, we expect δ ~ 90° for molecules lying in-plane (Fig 1). This value would increase to about δ ~ 110° for η = 70° and δ ~ 150° at η = 45° (Supplementary Fig. S1). The population with δ > 120° (η < 60°) observed in Fig. 3a is thus attributed to filaments tilted by more than 30° from the sample plane. Thus, selecting only delta values with δ < 110° (η > 70°, i.e. within 20° from the sample plane) allows to measure actin filament organization in SFs without a strong bias on σ_Δρ_ (Supplementary Fig. S5). Using this criterion, the parameter σ_Δρ_ can therefore be used as a minimally-biased quantitative estimate of in-plane actin filament organization in selected regions. σ_Δρ_ is seen to decrease in SFs for δ < 110° (Fig. 3e-f, ROIs 1 and 2, compare with Fig. 2f,h), reaching the σ_Δρ_ values in single actin filaments (Fig. 1 g), thus showing now highly aligned actin filaments in ventral SFs and FAs. Regions in dorsal SFs and meshworks depict now more clearly populations of actin filaments crossing each other, as expected (Fig. 3e-f, ROIs 3 and 4, compare with Fig. 2f) (see also Supplementary Fig. S7). Importantly, the estimated σ_Δρ_ is found to be of similar values independently of the radius size thresholding (data not shown), as long as in-plane filaments are selected (δ < 110°).

Our analysis emphasizes two novel possibilities that the 4polar-STORM method offers for the investigation of actin filament organization. First, the wobbling of single fluorophores can be used for the quantification of their off-plane tilt orientations, exploiting the degree of bias even at a slightly reduced detection NA. Second, the lowest wobbling values obtained in a population of fluorophores can be used to select in-plane populations of actin filaments and study their organization in a quantitative manner. Not quantifying these sources of bias can lead to errors of interpretation in the measured organizations, especially at the high detection NAs generally used^22,23,24^. In what follows, we exploit these considerations to quantitatively compare actin filament organization in different structures of the cell.

### Actin filament organization imaging by 4polar-STORM in-plane selection

We first analysed actin filament organization in different types of SFs in cultured U2OS cells on unpatterned or micropatterned coverslips (see Methods), quantifying σ_Δρ_ for molecules with δ < 110°. In what follows, the PSF radius of single molecules is not thresholded, results being very similar in both conditions. Regions defining ventral, peripheral, dorsal SFs as well as transverse arcs and FAs are measured by 4polar-STORM. We combine phalloidin stainings with immunostainings for a FA protein, the phosphorylated form of focal adhesion kinase, p-FAK, in order to define the different types of SFs based on their association with FAs^42,43^ (Fig. 2a). A z-stack of the cell further allows us to visualize off-plane tilted structures (Fig. 2a). While ventral, peripheral and arc SFs are lying in the sample plane, dorsal SFs are the most tilted ones. σ_Δρ_ values were measured in rectangular ROIs of typically (0.5-1 μm) x 100 nm in size. σ_Δρ_ exhibits very large distributions, due to variations in actin filament organization among SFs within a given cell and across cells (Fig. 4a). Among the measured SFs, ventral SFs exhibit the highest filament alignment (lowest σ_Δρ_ value), with the average σ_Δρ_ close to that measured in single filaments (<σ_Δρ_ > ~ 25°). σ_Δρ_ values for transverse arc SFs are slightly higher, suggesting the presence of less well-aligned actin filaments or/and filaments crossing each other. Such organization is fully consistent with the proposed mechanisms of transverse arc assembly, involving both the progressive fusion and alignment of actin filament fragments and myosin filament stacks, and connections of forming arcs with dorsal SFs^44,45,46^. σ_Δρ_ values for peripheral SFs are, surprisingly, even larger, suggesting that peripheral SFs contain a larger population of actin filaments in various directions. This observation correlated with the fact that the measured peripheral SFs were often thicker than the measured ventral ones, which suggests that thicker SFs contain more various actin filaments directions. This hypothesis is in line with observations of actin bundles fusing with or splitting from peripheral fibers and with recent work showing that peripheral SFs and the cortical meshwork form a continuous contractile network ^47^. FAs, regardless their association with ventral or dorsal SFs, depict a much wider distribution of σ_Δρ_ values, containing both highly aligned actin filaments and actin filaments in mixed orientations. We hypothesize that this organization reflects the dynamical nature of FAs, whose precise assembly and maturation depend on the local mechanical environment^48,49^. At last, dorsal SFs exhibit the highest σ_Δρ_ values with <σ_Δρ_ > ~ 35°. We attribute these high values to the very nature of dorsal SF assembly involving extensive interconnections along their length with transverse arc SFs^42,43^. Depicting all <δ> values measured in all ROIs confirms that peripheral SFs, FAs and dorsal SFs contain the largest population of filaments tilted off plane, with the lowest proportion of single molecules with δ < 110° (Fig. 4a).

**Figure 4.**
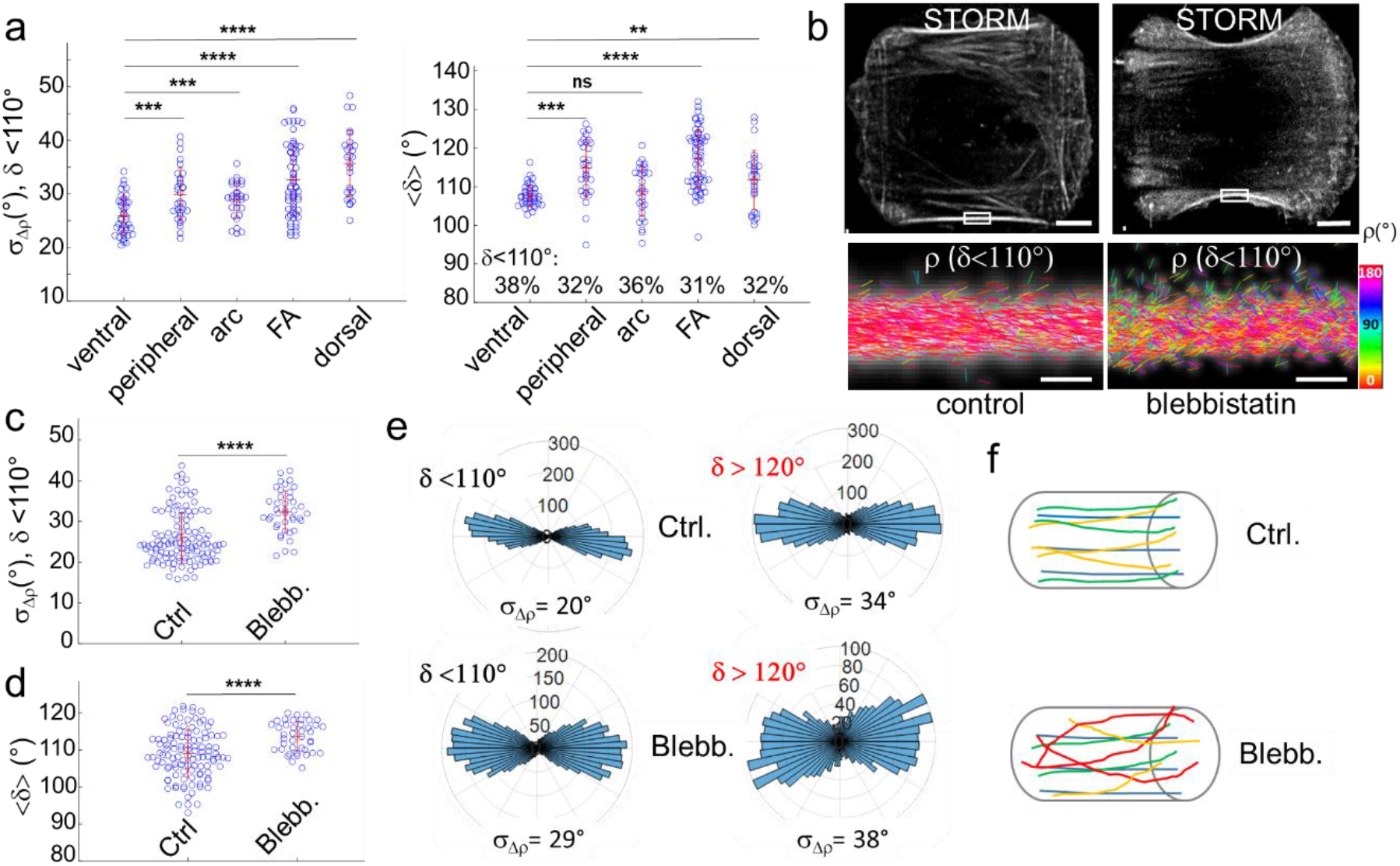
4polar-STORM imaging of actin filament organization in different types of stress fibers. (a) Left: σΔρ (standard deviation of Δρ = ρ-<ρ>) values measured in ROIs on different types of SFs, using only molecules with δ < 110°. The numbers of measured ROIs are as follows: n=42 (ventral), 32 (peripheral), 30 (arc), 51 (FA), 28 (dorsal) from a total of 8 cells. Typically, a few thousands of molecules per ROI are measured. Right: same analysis, plotting <δ> (average of δ measured for each ROI, all δ values selected). For each SF type, the percentage of the δ population with δ < 110° is indicated. Statistical significance of scatter plots: ns (p<0.05); ** (p < 0.01); *** (p < 0.001); **** (p < 0.0001) (T-test). All comparisons are made with respect to the ventral SF population. (b) top: STORM images of control and blebbistatin-treated U2OS cells on micropatterns. Scale bars, 6.5 μm; bottom: 4polar-STORM ρ stick images of zoomed regions (rectangles in the STORM images). Scale bars, 500 nm. (c) Effect of blebbistatin on peripheral SFs (see Methods for blebbistatin treatment). σ_Δρ_ values reported for all ROIs, using only molecules with δ < 110°. 4 cells were analysed per condition, using in total 87 ROIs in control cells and 43 ROIs in treated cells. (d) <δ> values reported for all ROIs (all δ selected). (e) Polar-plot histograms of ρ in the zooms shown in (b), for δ < 110° (left) and δ > 120° (right). The corresponding σ_Δρ_ values are shown. (f) Schematic representations of the SFs observed in control and blebbistatin-treated conditions, showing highly-aligned actin filaments in 2D (top) and disorganized filaments in 3D (bottom).

These results show that while peripheral SFs are known to be under a larger mechanical strain than central ventral SFs^50^, their higher mechanical tension is not necessarily accompanied with a higher alignment of actin filaments in-plane. To probe the sensitivity of actin filament organization in SFs to contractility, we treated U2OS cells with blebbistatin, a drug that inhibits myosin II activity and therefore acto-myosin bundle contractility. To minimize dispersions due to SF heterogeneities, we performed this experiment on cells adhering to H-shaped micropatterned substrates of a well-defined size, concentrating on peripheral SFs (see Methods). Blebbistatin treatment leads eventually to the dissociation of the contractile peripheral SFs^51^ (also seen in our data). Blebbistatin concentration and incubation time were thus kept low enough to induce a loss in contractility, evidenced by the concave shape of relaxed SFs, while preserving the apparent macroscopic integrity of the SFs (Fig. 4b). The single molecule localization image alone (STORM, Fig. 4b) cannot inform us on the underlying actin filament organization in the relaxed SFs. 4polar-STORM measurements, however, show a slight decrease of actin filament alignment, with a statistically significant increase of <σ_Δρ_ > from 26° to 32° (Fig. 4c), and also an enrichment of 3D oriented filaments as seen from the increase of <δ> (Fig. 4d). Contractility loss induced by myosin II inactivation is therefore correlated to a decrease of actin filament organization which is also visible from the 4polar-STORM ρ images (Fig. 4b). Representative distributions of single-molecule orientations ρ in control and blebbistatin-treated SFs are shown in Fig. 4b and quantified in Fig. 4e, for both in-plane (δ < 110°) and off-plane (δ > 120°) filament populations. The spreading of the distributions induced by blebbistatin confirms a decrease in actin filament alignment and a concomitant increase in 3D oriented filament populations as schematically represented in Fig. 4f. 4polar-STORM measurements thus reveal that actin filament organization in SFs is sensitive to acto-myosin contractility, with myosin II inhibition inducing a loss in actin filament alignment at the nanometer-scale.

At last, we investigated actin filament organization in dense networks at the leading edge of B16 melanoma cells (Fig. 5a). We concentrate in particular on the lamellipodium, which spans a few micrometers at the cell border^36^. Again, the single molecule localization images alone (STORM in Fig. 5a,d) do not provide any information on how single AF488-phalloidin molecules orient with respect to one another and thus on the nanometer-scale organization of the labelled actin filaments; the very high filament density in these networks makes it even more challenging to decipher the precise arrangement of filaments. Remarkably, the contour of the cells imaged by 4polar-STORM shows different (ρ,δ) distributions than in the rest of the cell (Fig. 5b,c). Within the first hundreds of nanometers from the cell contour, single molecules exhibit wide distributions of orientations (ρ) and high wobbling (δ) values, signatures of more disorganized, off-plane actin filament populations. Selecting only in-plane filaments (δ < 110°) shows that actin filaments in these dense networks are not oriented isotropically, but that there are visible preferred orientations which cannot be detected in the single molecule localization images alone^25^ (Fig. 5d and quantification in Fig. 5e). Single molecule orientation distributions at the cell border (ROIs 1,2,4-8 in Fig. 5b) reveal in general the presence of two main populations with different contributions (Fig. 5e). Generally, one of the two orientation populations appears to predominate (ROIs 1,6-8). However, in some regions (ROIs 5), the two populations tend to contribute equally into a bimodal distribution, which becomes more isotropic when off-plane molecules are considered (δ > 120°). This behaviour has been observed in all cells measured (see another example in Supplementary Fig. S8). We note that these distributions are very different from those found in SFs, which exhibit much narrower distributions with a preferred orientation, even when considering off-plane molecules (ROI 10). At last, narrow distributions are found in microspikes (ROIs 3,9), which is a signature of parallel actin bundles within the lamellipodium, despite their close proximity to bimodal distributions (see ROIs 2 and 4 which surround ROI 3).

**Figure 5.**
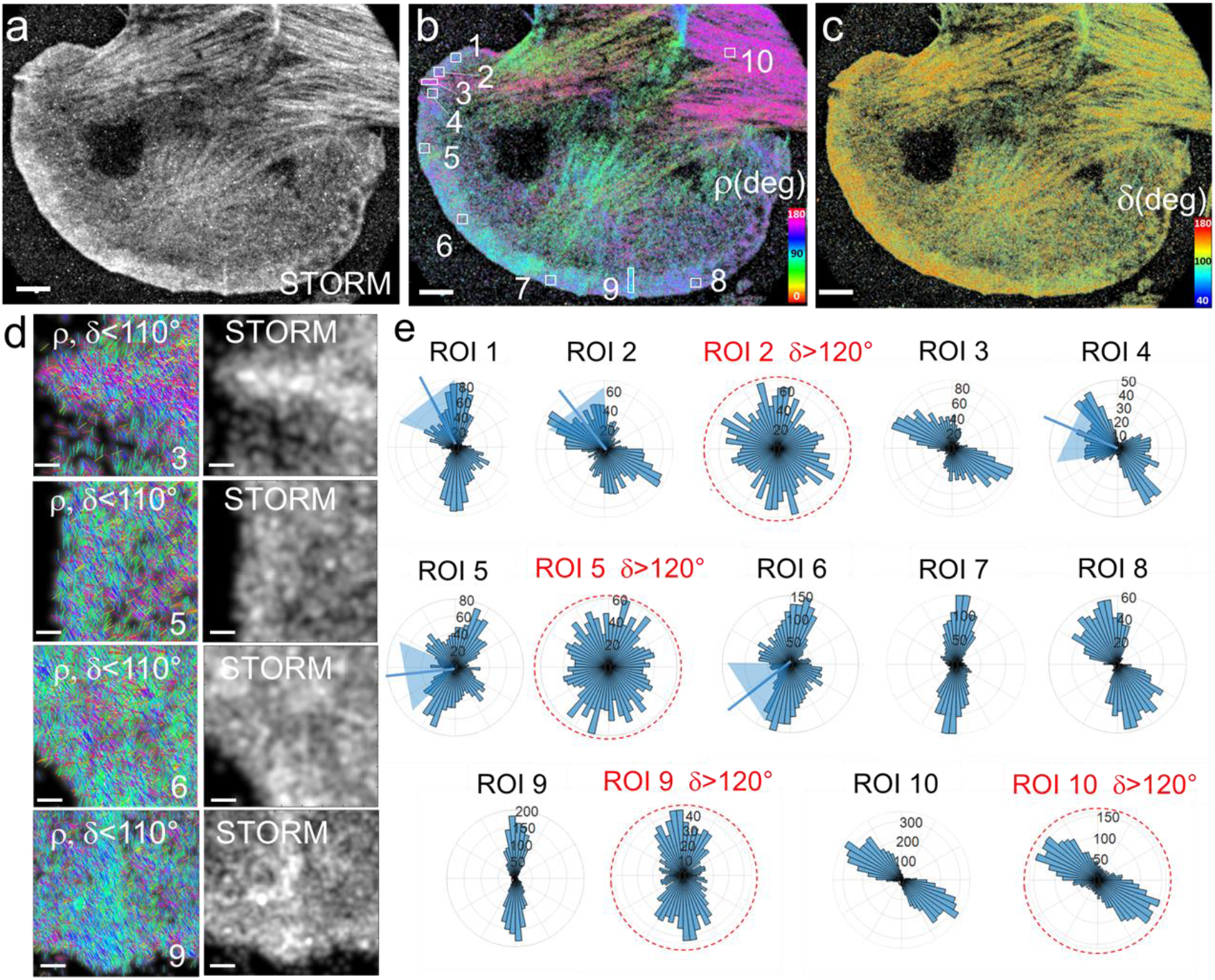
4polar-STORM imaging of actin filament organization in lamellipodia. (a) Single molecule localization STORM image of a B16 cell labelled with AF488-phalloidin. (b) Corresponding 4polar-STORM ρ stick image with color-coded orientation measurements. (c) 4polar-STORM δ stick image with color-coded wobbling angle measurements. (d) Examples of ρ stick images showing molecules with δ < 110° and corresponding STORM images in selected ROIs (squares in (b)). ROIs 1-9, regions in the lamellipodium; ROI 3,9, microspikes; ROI 10, SF. (e) Polar-plot histograms of ρ for the regions shown in (b). The condition δ < 110° is used, except for red-circled histograms for which δ > 120° molecules are selected. Scale bars (a-c), 4 μm; (d), 260 nm.

Remarkably, the angle between the two peaks of the observed bimodal distributions is close to 70°, with some variations along the cell contour, and points towards the normal to the membrane contour (Fig. 5e). This is reminiscent of observations made by EM where actin filaments display a bimodal angular distribution, with filament orientations peaking at 35° and −35° with respect to the direction of membrane protrusion^34,35,36^. This so-called dendritic organization was attributed to the angle imposed by the Arp2/3 complex involved in actin filament branching in those regions^36,52^. Bimodal orientation distributions in 4polar-STORM images were present in ROIs from hundreds of nanometers to micrometers sizes, and were variable depending on the region of the cell contour, emphasizing the importance of large field of view observations.

The variations in the precise distribution of actin filament orientations at the cell edge, and the non-negligible presence of 3D orientations, are consistent with recent findings in the literature based on EM and modelling^4^. A range of angular distributions of actin filament orientations has been evidenced, which appears to be more complex and heterogeneous than a pure bimodal distribution with peaks positioned at the 70° branching angle^53^. This angular spread depends on the cell mechanical load^54^, and the precise geometry of filament assemblies, not necessarily pointing towards the membrane normal, depends on the modulation of the protrusion rate^55,56^. Additionally, actin filament organization within the lamellipodium sheet is known to extend in 3D, with possibly different organizations at the cell surface and in the upper layer as evidenced recently in studies using 3D super-resolution localization microscopy^25^ or cryo-tomography EM^57^. The 4polar-STORM results, which do not differentiate between specific actin layers, suggest the co-existence of both preferred angular distributions of actin filament orientations and populations of 3D oriented filaments.

## Discussion

We developed a new super-resolution fluorescence microscopy method, called 4polar-STORM, which allows simultaneous measurements of single molecule localization and orientation in 2D, as well as an indirect evaluation of their 3D orientation. The 4polar-STORM analysis permits to evaluate quantitatively both the orientation and wobbling angles of single molecules, thus enabling the accurate determination of their organization. A detailed theoretical analysis of the dependence of the orientation and wobbling parameters on different sources of bias, either originating from SNR conditions or physical features, such as 3D molecular orientations or a high detection NA, is used to define appropriate detection conditions and analysis procedures for the minimally biased determination of molecular organization at the nanometer scale. It is important to note that the majority of polarized single molecule experiments have been performed under high NA conditions so far. This prevents the physical interpretation of wobbling values and thus an unbiased determination of the underlying molecular organization^23,24^. Other alternatives such as PSF engineering necessitate more complex signal processing to achieve quantitative measurements, and can be challenged in high density-labelled samples^14,19^.

To highlight the potential of 4polar-STORM to measure molecular organization in complex protein assemblies, we measured the nanometer-scale organization of actin filament-based structures involved in the adhesion and motility of mammalian cells. We focused on different types of stress fibers made of actin filament bundles and on the actin filament meshwork of the lamellipodium. We exploited the sensitivity of orientation and wobbling parameters to molecules lying off-plane to evidence the non-negligible contribution of 3D orientations in the measured populations of actin filaments, both in SF bundles and in the flat lamellipodium. Selecting the lowest wobbling values from measured single molecules permits to select only in-plane filament populations and determine their organization in a quantitative manner. This analysis permitted to evidence the very high actin filament alignment in all types of SFs, in line with EM studies^31,32^, but to also reveal differences in their nanometer-scale organization, consistently with their different modes of assembly and function in the cell. Thin 2D ventral and transverse arc SFs are made of highly aligned actin filaments, while thick peripheral SFs, off-plane oriented dorsal SFs, and FAs containing filament populations of mixed orientations, were seen to contain a non-negligible population of filaments with 3D off-plane orientations. Low doses of blebbistatin that inhibited myosin II activity in contractile peripheral SFs, while preserving their macroscopic integrity, resulted in a perturbation of the nanometer-scale organization of actin filaments, emphasizing the key role of myosin II in the organization of actin filaments in contractile SFs. Importantly, in-plane measurements of actin filament organization permitted us to investigate the organization of dense assemblies that is not accessible by single molecule localization imaging alone, in particular in the lamellipodium at the leading edge of motile cells. 4polar-STORM imaging revealed that the actin filaments in the lamellipodial meshwork are not oriented randomly but that they organize in preferred angular distributions, including bimodal distributions previously observed by electron microscopy. It is important to note that these characteristic features of the dendritic organization of the lamellipodium have not been observed with other super-resolution light microscopy methods.

These results show altogether the added value of combining localization and orientation measurements, and highlight the potential of 4polar-STORM to investigate the nanometer-scale organization of actin filaments in complex arrangements which are hardly accessible in standard optical super-resolution microscopy. In particular, the complex geometries of actin filaments close to the cell membrane and their link with local contractile and protrusive activity are poorly understood. Additionally, little is known on the organization of actin filaments in cell adhesion-mediating structures which play a large role in cell mechanics, in particular focal adhesions. Assessing the nanometric organization of actin filaments, and the contribution of 2D vs 3D oriented populations of filaments, at various stages of FA formation and maturation promises to permit us to understand how such complex molecular machineries assemble in order to sense, respond and adapt to mechanical stimuli. We note that while AF488-phalloidin is used in the present study, our method is fully compatible with a wide variety of labels as long as the orientational flexibility of the fluorophore is not very high. In particular, actin labelling by Atto633-phalloidin^10^ or silicon rhodamine-jasplakinolide (SiR-actin)^24,58^, have been reported to exhibit low wobbling. Genetically encoded probes for actin filament orientation studies are also currently being developed^59^, using strategies which could be extended to other proteins of interest. Such approaches which are amenable to live-cell measurements promise to open new directions for orientational dynamics studies, combining for instance single molecule orientation measurements with single particle tracking PALM^60^. Wobbling measurements in the case of fluorescent proteins, whose size is typically comparable to the size of the labelled protein of interest, might also reveal packing-related constraints and could thus provide additional protein organization readouts. Although the present work focuses on the implementation of 4polar-STORM for deciphering the organization of actin filaments, the framework we built regarding theoretical considerations for the retrieval of orientation parameters and the definition of appropriate acquisition and analysis procedures, can be easily adapted for the study of any molecule. The methodology of 4polar-STORM is also compatible with two-color localization and orientation measurements and thus can provide insights into the functional interplay between the nanometric organizations of interacting partners; the link between conformational changes in activated integrins and actin filament remodeling is such an example. At last, 4polar-STORM is compatible with 3D localization schemes, including astigmatism^61^ or multiplane^62,63^ strategies, and can therefore be adapted for exploring the full 3D organization of a large variety of biological structures^64^.

## Materials and Methods

### 4polar-STORM optical setup

Measurements are carried-out on a custom epi/TIRF-fluorescence microscope, whose detection path is adapted to retrieve four polarization states of the single molecule fluorescence images. The excitation light source is a continuous laser emitting at 488 nm (Sapphire 488LP-200, Coherent), whose beam is expanded by a telescope and circularly polarized by a quarter waveplate (AQWP10M-580, Thorlabs). A set of mirrors reflect the beam towards the microscope, followed by a large focal length lens (f = 400 mm) to focus the beam in the back focal plane of the objective, to provide an illumination field of view with a diameter of about 100 μm. After the reflection on a dichroic mirror (DI02-R488, Semrock Rochester NY), the excitation light is focused onto the sample by an oil immersion objective lens (Plan Apo 100×, NA = 1.45, Nikon). The emitted fluorescence is collected back by the same objective lens, passes through the dichroic mirror and a band pass emission filter (FF01-525/40, Semrock Rochester NY). At the microscope exit a non-polarizing beam splitter separates the beam in two paths, each of them being built up with a 1x relay imaging telescope that uses two (f 150 mm) lenses. In the first path, a Wollaston prism (separation angle 5°, CVI Laser Optics) is placed at the back focal intermediate image plane, aligned for 0°-90° polarization split. In the second path, a similar Wollaston prism is placed just after an achromatic half wave plate (AHWP05M-600, Thorlabs), to provide 45°-135° polarized images. The two beams are recombined by a mirror reflection of the first path, and refocused on the EMCCD camera detection plane (iXon Ultra 888, Andor, 13 μm pixel size), such as to fill the CCD chip with four polarized images. The size of the images is set by a diaphragm placed in the first image plane at the exit of the microscope (typical image field of view, 40 μm x 40 μm). In addition, two diaphragms are placed in intermediate planes conjugated to the back focal plane of the objective in order to reduce the detection numerical aperture to NA_det_ = 1.2. The imaging lens provides a total magnification of ×100, corresponding to a pixel size of 130 nm on the EMCCD. The stability of the focus throughout the measurement is ensured by a commercial system (Perfect Focus System, Nikon). For initial positionning, the sample is mounted on a XYZ piezo stage (Physik Instrumente). The acquisition parameters are controlled by a commercial imaging software (AndorSolis, Andor). For STORM imaging, a first fluorescence image is recorded with low intensity (~ 500 W/cm^2^, below STORM blinking conditions), ensuring the identification of relevant parts of the sample. The intensity is then raised to 5-8 kW/cm^2^, which is a typical level to provide a good compromise between signal level and blinking rate. The images are acquired at a rate of 100 ms/image, camera gain 300, with a total of about 30,000 images depending on the molecular density.

### Cell culture

4polar-STORM measurements were made in U2OS osteosarcoma cells (Fig. 1–4) and B16-F1 mouse melanoma cells (Fig. 5). Naive U2OS cells (gift from Flavio Maina, IBDM, France) were used for assessing the effect of blebbistatin. U2OS CA-MLCK cells (gift from Sanjay Kumar, UC Berkeley, USA) cultured in 0 ng/mL doxycycline were used for all other experiments. U2OS cells were maintained in McCoy’s 5A medium (ThermoFisher Scientific, 26600-080) supplemented with 10% fetal bovine serum (Biowest, S181H), 100 U/mL penicillin and 100 μg/mL streptomycin (Sigma, P4333) in a humidified incubator at 37°C and 5% CO2. B16-F1 cells (gift from Klemens Rottner, Technische Universität Braunschweig, Germany) were cultured in DMEM (ThermoFisher Scientific, 41966-029) supplemented with 10% fetal bovine serum (PAA Laboratories, A15-102), 100 U/mL penicillin and 100 μg/mL streptomycin (Sigma, P4333) in a humidified incubator at 37°C and 5% CO2.

### Cell preparation for 4polar-STORM. U2OS cells

24 mm-diameter high-precision (170 μm ± 5 μm) glass coverslips (Marienfeld, 0117640) were cleaned with base piranha (Milli-Q water, 30% ammonium hydroxide, 35% hydrogen peroxide at a 5:1:1 volume ratio) for 15 min, rinsed with Milli-Q water for 2 x 5 min in a bath sonicator, sonicated in 70% ethanol for 5 min, and air-dried before coating with fibronectin (SIGMA F1141) for 2 h at room temperature (RT) and at a final fibronectin concentration of 20 μg/mL in PBS. For experiments with micropatterned substrates, medium-size (1100 μm^2^) H-shaped patterns from CYTOO (10-900-00-18) were similarly coated with 20 μg/mL fibronectin. U2OS cells were seeded onto fibronectin-coated coverslips and allowed to spread for 5 h on micropatterned substrates or overnight on nonpatterned ones. Cells were fixed for 15 min with 4% formaldehyde (Electron Microscopy Sciences 15714) in cytoskeleton buffer (10 mM MES pH 6.1, 150 mM NaCl, 5 mM EGTA, 5 mM MgCl_2_, 5 mM glucose), washed for 2 x 5 min in PBS, then permeabilized and blocked in phosphate-buffered saline (PBS) containing 0.1% saponin and 10% bovine serum albumin (BSA) for 1 h at RT. Cells were incubated successively with primary rabbit anti-phospho-FAK antibodies at 1:200 (ThermoFisher Scientific 44-624G) and secondary donkey anti-rabbit Alexa Fluor 647-conjugated IgG secondary antibodies at 1:1000 (ThermoFisher Scientific A-31573) each for 1 h at RT and with three 10-min washes in-between antibody incubations. After five 6-min washes, cells were incubated with 0.5 μM Alexa Fluor 488 (AF488)-phalloidin (ThermoFisher Scientific A12379) in 0.1% saponin/10% BSA/PBS overnight at 4°C in a humidified chamber. For 4polar-STORM measurements, coverslips with stained cells were mounted in an Attofluor cell chamber (ThermoFisher Scientific A7816) with freshly prepared STORM imaging buffer (see composition below) and the chamber covered with a glass coverslip to minimize contact with oxygen. To visualize focal adhesions in order to define the types of stress fibres measured, AF488-phalloidin and phospho-FAK-co-stained cells were imaged before each STORM acquisition on an optical setup employing a confocal spinning disk unit (CSU-X1-M1 from Yokogawa) connected to the side-port of an inverted microscope (Eclipse Ti-U from Nikon Instruments), using a Nikon Plan Apo ×100/1.45 NA oil immersion objective lens, 488- and 641-nm laser lines (Coherent) and an iXon Ultra 888 EMCCD camera (1024×1024 pixels, 13×13 μm pixel size, Andor, Oxford Instruments). z-stacks were acquired with a Δz interval of 0.5 μm.

### B16-F1 cells

24 mm-diameter high-precision (170 μm ± 5 μm) glass coverslips (Marienfeld, 0117640) were sonicated in 70% ethanol for 5 min and air-dried before coating with mouse laminin (SIGMA L2020) for 1 h at RT and at a final laminin concentration of 25 μg/mL in coating buffer (50 mM Tris-HCl pH 8, 150 mM NaCl). B16-F1 cells were seeded onto laminin-coated coverslips and allowed to spread overnight. To stimulate lamellipodia formation, cells were treated with aluminum fluoride for 15 min by adding AlCl_3_ and NaF to final concentrations of 50 μM and 30 mM, respectively, in pre-warmed, full growth medium. Cells were fixed for 20 min with a mixture of prewarmed (37°C) 0.25% glutaraldehyde (Electron Microscopy Sciences 16220) and 4% formaldehyde (Electron Microscopy Sciences 15714) in cytoskeleton buffer, and treated with fresh sodium borohydride (1 mg/mL) in PBS for 2 × 5 min to reduce background fluorescence. Cells were washed in PBS for 3 × 5 min before an overnight incubation with 0.5 μM AF488-phalloidin in 0.1% saponin/10% BSA/PBS at 4°C in a humidified chamber. For 4polar-STORM measurements, coverslips were mounted as for U2OS cells.

### Blebbistatin treatment

Blebbistatin from Sigma (B0560) was prepared at 10 mM in DMSO. U2OS cells were seeded onto fibronectin-coated medium-size H-shaped patterns from CYTOO and allowed to spread for 5 h, as detailed above. Cells were incubated for 15 min with 50 μM blebbistatin (i.e. in medium also containing 0.5% DMSO due to the blebbistatin stock dilution), or with medium containing 0.5% DMSO (control cells). Cells were fixed with 4% formaldehyde in cytoskeleton buffer for 15 min, and washed in PBS for 2 x 5 min before an overnight incubation with 0.5 μM AF488-phalloidin in 0.1% saponin/10% BSA/PBS at 4°C in a humidified chamber. Cells were mounted for 4polar-STORM measurements as detailed above.

### Reconstitution of single actin filaments for 4polar-STORM

Lyophilized rabbit skeletal muscle G-actin (Cytoskeleton, Inc. AKL99) was resuspended to 5 mg/mL (119 μM) in G-buffer (5 mM Tris-HCl pH 8, 0.2 mM Na_2_ATP, 0.1 mM CaCl_2_, 1 mM DTT), aliquots snap-frozen in liquid nitrogen and stored at −80°C. Frozen aliquots were thawed and centrifuged for 30 min at 120,000 g in a benchtop Beckman air-driven ultracentrifuge (Beckman Coulter Airfuge, 340401) to clear the solution from aggregates. Clarified G-actin was kept at 4°C and used within 3-4 weeks. For reconstitution experiments, G-actin was polymerized at 5 μM final concentration in actin polymerization buffer (5 mM Tris-HCl pH 8, 50 mM KCl, 1 mM MgCl_2_, 0.2 mM Na_2_ATP, 1 mM DTT) in the presence of 5 μM AF488-phalloidin for at least 2 h at RT. Flow cells for measurements on reconstituted actin filaments were prepared as follows. Microscope glass slides and coverslips were cleaned for 15 min in base-piranha solution, rinsed twice, 5 min each, with Milli-Q water in a bath sonicator, and stored in ethanol up to one month. To assemble flow cells, slides and coverslips were dried with synthetic air, and ~10 μL channels were assembled by sandwiching ~2-mm-wide and ~2.5-cm-long strips of Parafilm between a cleaned glass slide and coverslip and melting on a hot plate at 120°C. The chambers were incubated for 45 min with 1 M KOH, rinsed with actin polymerization buffer, incubated for another 15 min with 1 mg/mL poly-L-lysine (PLL; Sigma P8920), and rinsed with actin polymerization buffer. Reconstituted AF488-phalloidin-labelled actin filaments were diluted to 0.1-0.2 μM, loaded into the PLL-coated flow channels and left for 15 min to immobilize actin filaments. Actin polymerization buffer was then exchanged with STORM imaging buffer (see composition below), and flow channels sealed with VALAP (1:1:1 vasoline:lanoline:paraffin). The typical experimental conditions were TIRF illumination, laser power 150 mW, camera gain 300 and 200-ms integration time. A stack of 5000 images was used for 4polar-STORM imaging. The materials and chemicals for glass cleaning were as follows. Glass slides (26×76 mm) from Thermo Scientific (AA00000102E01FST20). Glass coverslips (24×60 mm) from Thermo Scientific (BB02400600A113FST0). Ammonium hydroxide solution from SIGMA (221228). Hydrogen peroxide solution from SIGMA (95299).

### STORM imaging buffer preparation

The final composition of the buffer for 4polar-STORM measurements was 100 mM Tris-HCl pH 8, 10% w/v glucose, 5 U/mL pyranose oxidase (POD), 400 U/mL catalase, 50 mM β-mercaptoethylamine (β-MEA), 1 mM ascorbic acid, 1 mM methyl viologen, and 2 mM cyclooctatetraene (COT). D-(+)-glucose was from Fisher Chemical (G/0500/60). POD was from Sigma (P4234-250UN), bovine liver catalase from Calbiochem/Merck Millipore (219001-5MU), β-MEA from Sigma (30070), L-ascorbic acid from Sigma (A7506), methyl viologen from Sigma (856177), and COT from Sigma (138924). Glucose was stored as a 40% w/v solution at 4°C. POD was dissolved in GOD buffer (24 mM PIPES pH 6.8, 4 mM MgCl_2_, 2 mM EGTA) to yield 400 U/mL, and an equal volume of glycerol was added to yield a final 200 U/mL in 1:1 glycerol:GOD buffer; aliquots were stored at − 20°C. Catalase was dissolved in GOD buffer to yield 10 mg/mL, and an equal volume of glycerol was added to yield a final 5 mg/mL (230 U/μL) of catalase in 1:1 glycerol:GOD buffer; aliquots were stored at −20°C. β-MEA was stored as ~77 mg powder aliquots at −20°C; right before use, an aliquot was dissolved with the appropriate amount of 360 mM HCl to yield a 1 M β-MEA solution. Ascorbic acid was always prepared right before use at 100 mM in water. Methyl viologen was stored as a 500 mM solution in water at 4°C. COT was prepared at 200 mM in DMSO and aliquots stored at −20°C. After mixing all components to yield the final buffer composition, the buffer was clarified by centrifugation for 2 min at 16,100 g, and the supernatant kept on ice for 15 min before use. Freshly prepared STORM buffer was typically used within a day.

## Supplementary Information

Supplementary Note 1. Model and retrieval of orientation parameters

Supplementary Note 2. Calibration factors in 4polar-STORM

Supplementary Note 3. Data processing algorithm of the 4polar-STORM method Supplementary Figure S1. Retrieval bias on δ

Supplementary Figure S2. Cone model used for orientation parameter retrieval Supplementary Figure S3. Camera noise estimation.

Supplementary Figure S4. 4polar-STORM d images of F-actin in cells Supplementary Figure S5. Retrieval bias on σ_Δρ_

Supplementary Figure S6. Statistics on detection parameters in 4polar-STORM imaging of F-actin in stress fibers in cells

Supplementary Figure S7. 4polar-STORM imaging of of F-actin in cells, selecting in-plane actin filament populations

Supplementary Figure S8. 4polar-STORM imaging of actin filament organization in lamellipodia.

## Acknowledgements

This research has received funding from the « Investissements d’Avenir » French Government program managed by the French National Research Agency ANR (ANR-16-CONV-0001), from Excellence Initiative of Aix-Marseille University A*MIDEX (ANR-11-IDEX-0001), European Union’s Horizon 2020 research and innovation programme under the Marie Sklodowska-Curie grant agreement No 713750, and the SEPTIMORF and 3DPolariSR ANR grants (ANR-17-CE13-0014; ANR-20-CE42-0003). This work was also supported by INRIA in the frame of NAVISCOPE-IPL (INRIA Project Lab). The authors are grateful to Arita Silapetere (Humboldt Universitat zu Berlin, Germany) for the assistance in the initiation of the project and Louwrens Van Dellen (Purplecode Aix en Provence, France) for help in the experimental acquisition programming. The authors further acknowledge useful discussions of the results with Klemens Rottner (Technische Universität Braunschweig, Germany).

## Author Contributions

S.B and M.M conceived and initiated the project. C.V.R built the optical system, performed experiments on cells and analysed the data with S.B. V.C performed experiments on single actin filaments and analysed the data. C.A.V.C wrote the data processing algorithm. C.V.R and S.B wrote part of the data analysis algorithms. All authors wrote the paper and contributed to the scientific discussion.

## Author information

The authors declare that they have no competing financial interests. Correspondence and requests for materials should be addressed to the corresponding author.

## Data availability

The datasets generated and analysed during the current study are available from the corresponding author on reasonable request.

## Code availability

The code used to process and analyse the data during the current study are available from the corresponding author on reasonable request.

