## Supplementary Notes and Figures for "4polar-STORM polarized super-resolution imaging of actin filament organization in cells"

**Content**

**Supplementary Note 1.** Model and retrieval of orientation parameters
**Supplementary Note 2.** Calibration factors in 4polar-STORM
**Supplementary Note 3.** Data processing algorithm of the 4polar-STORM method
**Supplementary Figure S1.** Retrieval bias on $\delta$
**Supplementary Figure S2.** Cone model used for orientation parameter retrieval
**Supplementary Figure S3.** Camera noise estimation
**Supplementary Figure S4.** 4polar-STORM $\delta$ images of F-actin in cells
**Supplementary Figure S5.** Retrieval bias on $\sigma_{\Delta\rho}$
**Supplementary Figure S6.** Statistics on detection parameters in 4polar-STORM imaging of F-actin in stress fibers in cells
**Supplementary Figure S7.** 4polar-STORM imaging of F-actin in cells, selecting in-plane actin filament populations
**Supplementary Figure S8.** 4polar-STORM imaging of actin filament organization in lamellipodia

### Supplementary Note 1. Model and retrieval of orientation parameters

A single molecule orientation is determined by its absorption dipole $\vec{\mu}_a$ and emission dipole $\vec{\mu}_e$, which lie along transition dipole moment directions for the respective absorption and emission transitions. In what follows, we suppose that the absorption and emission dipoles have the same orientation. This assumption is appropriate in the present work which focuses on the emission detection only, but different situations can also be derived from the following equations.

Dipole orientations $(\theta, \varphi)$ are defined in the frame of the distribution angles that the molecule explores during the integration time of the image (typically tens to hundreds ms, which is much longer than the rotational time of the molecules). We consider a molecule wobbling within a distribution $f(\theta, \varphi)$ during the integration time of the detector. $f(\theta, \varphi)$ is defined here as a cone function, of value 1 for $(0 \leq \theta \leq \delta/2)$ and 0 elsewhere (a Gaussian function would lead to very similar results). $\rho$ is the orientation of the projection of the cone in the sample plane $(x, y)$, and $\eta$ is the out-of-plane orientation of the cone, relative to $z$ (see Figure).
The absorption and emission dipoles $\vec{\mu}_a(\theta, \varphi)$ and $\vec{\mu}_e(\theta, \varphi)$ are therefore expressed in the macroscopic frame $(x, y, z)$ of the sample as:

$$\vec{\mu}_{a,e}(\theta, \varphi, \rho, \eta) = \mu_{a,e} \begin{bmatrix} \sin^2\rho + \cos\eta\cos^2\rho & (\cos\eta - 1)\sin\rho\cos\rho & \sin\eta\cos\rho \\ (\cos\eta - 1)\sin\rho\cos\rho & \cos^2\rho + \cos\eta\sin^2\rho & \sin\eta\sin\rho \\ -\sin\eta\cos\rho & -\sin\eta\sin\rho & \cos\eta \end{bmatrix} \begin{bmatrix} \sin\theta\cos\varphi \\ \sin\theta\sin\varphi \\ \cos\theta \end{bmatrix}$$

Eq. S1

Where $\mu_{a,e}$ is the dipole amplitude.

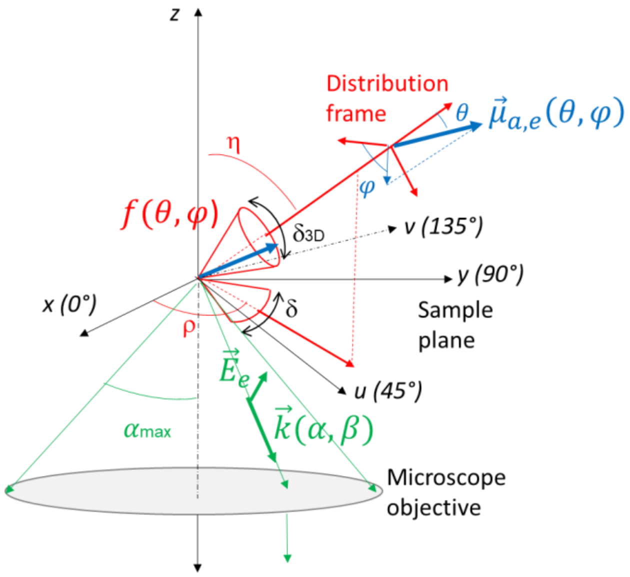


The fluorescence signal is the result of an excitation step, whose efficiency is quantified by the absorption probability, and an emission step, in which the emission dipole radiates. A complete derivation of the fluorescence signal from wobbling dipoles can be found for instance in[1]. Here we provide the expressions used in this work to extract the orientation parameters from single dipoles, from polarized intensities measured in the 4polar-STORM method.

**Excitation.** The absorption probability is proportional to[2]:

$$P_{abs}(\theta,\varphi,\rho,\eta) \propto \left|\vec{\mu}_a(\theta,\varphi,\rho,\eta)\cdot\vec{E}\right|^2 \quad \text{Eq. S2}$$

with $\vec{E}$ the excitation field.

In the case of normal incidence circular polarization, $\vec{E} = E_0(1,1,0)$ with $E_0$ the field amplitude.

In the case of total internal reflection (TIRF) illumination, $\vec{E} = (E_x, E_y, E_z)$ takes a more complex form that can be found in[3], which involves a contribution along $z$. In the present work, the incident polarization in TIRF is set such as the in-plane $(E_x, E_y)$ contributions are balanced, in order to minimize any in-plane photoselection.

**Emission.** The emission field radiated from the emission dipole $\vec{\mu}_e$ along the propagation direction $\vec{k}$ writes, in free space:

$$\vec{E}_e(\theta,\varphi,\rho,\eta,\vec{k}) \propto (\vec{k}\times\vec{\mu}_e(\theta,\varphi,\rho,\eta))\times\vec{k} \quad \text{Eq. S3}$$

In reality the dipole is placed within a medium of refractive index supposedly close to water, above an interface with glass. To account for these interfaces, expressions derived in[4] and[5] are used.

**Fluorescence intensity**. The fluorescence intensity detected along a polarization direction $\vec{\varepsilon}$ and a propagation vector $\vec{k}$ (see Figure) is deduced from the absorption and emission probabilities product[2,6,7]:

$$I_\varepsilon(\vec{k}) \propto \int_0^{2\pi} d\varphi \int_0^{\delta/2} d\theta\ \sin\theta\, P_{abs}(\theta,\varphi)\left|\vec{E}_e(\theta,\varphi,\rho,\eta,\vec{k})\cdot\vec{\varepsilon}\right|^2 \quad \text{Eq. S4}$$

Where $\propto$ encompasses all collection/excitation efficiency factors that do not affect the present analysis. To account for the detection numerical aperture (NA), it is necessary to integrate all vector fields directions $\vec{k}(\alpha,\beta)$ over the collected NA, with $(0 \le \alpha \le \alpha_{max}, 0 \le \beta \le 2\pi)$. The total detected intensity writes therefore as:

$$I_\varepsilon = \int_0^{2\pi} d\beta \int_0^{\alpha_{max}} d\alpha\ \sin\alpha\, I_\varepsilon(\vec{k}(\alpha,\beta)) \quad \text{Eq. S5}$$

Which means that at high NA integration, the detection is affected by a mixture of polarization contributions.

A simplified writing of this integration is[8]:

$$I_\varepsilon = \kappa_1 I_\varepsilon + \kappa_2 I_{\varepsilon_\perp} + \kappa_3 I_z$$

with $(\vec{\varepsilon},\vec{\varepsilon}_\perp,\vec{z})$ forming a direct orthonormal basis (in this work, either $(x,y,z)$ or $(u,v,z)$) and with the $(\kappa_1,\kappa_2,\kappa_3)$ coefficients follow a specific dependence on the NA [8]:

$$\kappa_3 = \frac{1}{3}(2 - 3\cos\alpha_{max} + \cos^3\alpha_{max})$$
$$\kappa_2 = \frac{1}{12}(1 - 3\cos\alpha_{max} + 3\cos^2\alpha_{max} - \cos^3\alpha_{max})$$
$$\kappa_1 = \frac{1}{4}(5 - 3\cos\alpha_{max} - \cos^2\alpha_{max} - \cos^3\alpha_{max})$$

In this work, the detection is performed along the polarization directions (0°, 90°, 45°, 135°), e.g. $\vec{\varepsilon} = \vec{x}, \vec{y}, \vec{u}, \vec{v}$ with $\vec{u} = (\vec{x} + \vec{y})/\sqrt{2}$ and $\vec{v} = (\vec{y} - \vec{x})/\sqrt{2}$. Equations S4, S5 show that there is a matrix relation between the measured intensities $(I_0, I_{90}, I_{45}, I_{135})$ and the dipole-orientation dependent quantities:

$$\langle \mu_{e,\varepsilon} \cdot \mu^*_{e,\varepsilon'} \rangle(\rho, \delta, \eta) = \int_0^{2\pi} d\varphi \int_0^{\delta/2} d\theta \ \sin\theta \, P_{abs}(\theta, \varphi) \, \mu_{e,\varepsilon}(\theta, \varphi, \rho, \eta) \cdot \mu^*_{e,\varepsilon'}(\theta, \varphi, \rho, \eta)$$

Eq. S6

with $(\varepsilon, \varepsilon')$= $(x, y, z)$.

To measure the molecular orientation parameters $(\rho, \delta)$ with the condition $(\eta = 90°)$, two independent measurements are required. Note that $\eta$ cannot be extracted from a 4polar-STORM measurement unless specific defocusing or interface-imaging conditions are realized which involve PSF fitting[1,9]. The present work is based on ratiometric intensity imaging, which does not require any PSF fitting. We define the two polarization factors, normalized with respect to the total intensity:

$$P_0 = \frac{I_0 - I_{90}}{I_0 + I_{90}} \qquad \text{and} \qquad P_{45} = \frac{I_{45} - I_{135}}{I_{45} + I_{135}}$$

Eq. S7

We denote the total intensity $I_T = I_0 + I_{90} + I_{45} + I_{135}$. Note that $I_0 + I_{90} = I_{45} + I_{135}$ in an ideal optical system.

From the measurements of the intensities $(I_0, I_{90}, I_{45}, I_{135})$ or the ratios $(P_0, P_{45})$, components $\langle \mu_{e,\varepsilon} \cdot \mu^*_{e,\varepsilon'} \rangle(\rho, \delta)$ with $(\varepsilon, \varepsilon') = (x, x), (y, y), (x, y)$ can be retrieved. These quantities depend on the orientation parameters $(\rho, \delta)$. Note that this measurement is an in-plane projection of polarizations, and is therefore insensitive to components involving $z$, e.g. the 3D orientation $\eta$ of the emission dipoles.

A very simple expression of the moments expressed in Eq. S6 can be given ignoring the 3D expansion of the cone distribution (e.g. $f(\theta, \varphi)$ is a 2D-flat cone $(\theta \in [0, \delta/2], \varphi \in [0, 2\pi])$. Supposing no photoselection along the in-plane excitation directions, which is achievable experimentally, this dependence is:

$$\langle \mu_{e,x} \cdot \mu^*_{e,x} \rangle(\rho, \delta) = \frac{\mu_e^2}{2}(\cos 2\rho \, \mathrm{sinc}\, \delta + 1)$$
$$\langle \mu_{e,y} \cdot \mu^*_{e,y} \rangle(\rho, \delta) = \frac{\mu_e^2}{2}(1 - \cos 2\rho \, \mathrm{sinc}\, \delta)$$
$$\langle \mu_{e,x} \cdot \mu^*_{e,y} \rangle(\rho, \delta) = \mu_e^2(\sin 2\rho \, \mathrm{sinc}\, \delta)$$

Eq. S8

The $(\rho, \delta)$ parameters can easily be extracted from Eq. S8.

Inspired from this derivation, a simplified procedure is developed to analyse 4polar-STORM images. The $(\rho, \delta)$ parameters retrieval is made under the assumption that the molecular wobbling distribution lies within the sample plane with $(\eta = 90°)$. We consider also that there is no specific photoselection between the $x$ and $y$ (a $z$ contribution of the excitation field in the absorption probability can be possibly introduced if using TIRF). Under the paraxial approximation for the detection, the fluorescence intensity along the detection polarization, for an emission dipole $\vec{\mu}_e(\theta, \varphi)$ oriented along $(\theta, \varphi)$, can then be simplified into:

$$I_\varepsilon(\theta, \varphi, \rho, \eta) \propto |\vec{\mu}_e(\theta, \varphi, \rho, \eta) \cdot \vec{\varepsilon}|^2 \qquad \text{Eq. S9}$$

with $\vec{\varepsilon} = \vec{x}\ (I_0), \vec{y}\ (I_{90}), \vec{u}\ (I_{45}), \vec{v}\ (I_{135})$, with $\vec{u} = (\vec{x} + \vec{y})/\sqrt{2}$ and $\vec{v} = (\vec{y} - \vec{x})/\sqrt{2}$.
Simplifying the cone distribution function into a 2D-flat cone $(\theta \in [0, \delta/2], \varphi \in [0, 2\pi])$, the integration over the distribution function simplifies the detected intensities into :

$$I_0 = \frac{I_T}{2}(\cos 2\rho \operatorname{sinc}\delta + 1) \text{ and } I_{90} = \frac{I_T}{2}(1 - \cos 2\rho \operatorname{sinc}\delta)$$
$$I_{45} = \frac{I_T}{2}(\sin 2\rho \operatorname{sinc}\delta + 1) \text{ and } I_{135} = \frac{I_T}{2}(1 - \sin 2\rho \operatorname{sinc}\delta) \qquad \text{Eq. S10}$$

With $\operatorname{sinc}\delta = \frac{\sin\delta}{\delta}$ and $I_T$ the total intensity.
Under these conditions,

$$P_0(\rho, \delta) = \cos 2\rho \operatorname{sinc}\delta$$
$$P_{45}(\rho, \delta) = \sin 2\rho \operatorname{sinc}\delta \qquad \text{Eq. S11}$$

Therefore the orientation parameters can be easily deduced with

$$\rho = \frac{1}{2} \operatorname{atan}\left(\frac{P_{45}}{P_0}\right)$$
$$\operatorname{sinc}\delta = \sqrt{P_0^2 + P_{45}^2} \qquad \text{Eq. S12}$$

These expressions will be used to analyse the 4polar-STORM results, and their validity in real situations (high detection NA, tilted wobbling distribution) is discussed below. Importantly, Eq. S12 shows that the 4polar-STORM method permits to estimate $\rho$ and $\delta$ independently, which is not the case of polarization schemes that use only two polarization projections[10].

Equation S12 is valid for wobbling molecules lying in a flat cone in the sample plane and in the paraxial approximation. When molecular distributions resemble a full cone tilted off-plane $(\eta < 90°)$, the determined $\rho$ parameter is expected to be unchanged, however $\delta$ will be overestimated. This overestimation is expected to be even more dramatic when $\eta$ decreases away from 90° and when working at high NA, since the emission of highly tilted dipoles is mostly manifested at high NA, e.g. highly tilted $\vec{k}$ vectors[11]. The bias between the expected $\delta$ (denoted $\delta_{3D}$) and the measured $\delta$

(extracted from Eq. S12) is represented in Fig. S1 for different tilt angles $\eta$ of the cone distribution representing the molecular wobbling. To calculate this bias, expressions of the polarized intensities $(I_0, I_{90}, I_{45}, I_{135})$ are extracted from Eq. S5 using the TIRF illumination condition used in this work, and generating different sets of calculated $(\eta, \delta)$ parameters, supposing $\rho = 0°$ (this parameter does not influence the results). From these intensities, $\delta$ is determined using Eq. S12. The results show that at the working condition NA = 1.45, a strong bias on $\delta$ is visible even in wobbling cones lying in the sample plane ($\eta = 90°$). This bias is however strongly reduced when reducing the detection NA, which reduces the detection efficiency of tilted dipoles. At NA = 1.2, the bias on $\delta$ stays at reasonable level for δ values measured in this work (δ ~ 90°), as long as $\eta > 45°$ while keeping a reasonable signal to noise condition (about 60% of the total intensity is preserved for cones lying close to the sample plane with $\eta > 45°$). In order to provide an unbiased picture of δ values for in-plane lying wobbling cones, it is thus favourable to work at a detection NA close to 1.2.

The use of the 'flat cone' model used in Eq. S12 is shown to allow a close to unbiased determination of $\delta$ when the cone is lying in the sample plane (Fig. S1). The reason for this is that molecules are essentially excited in the sample plane, which decreases the weight of molecules possibly oriented off-plane in the final polarized intensities. The difference between the 'flat cone' model used in Eq. S12 and the full cone model has been assessed more precisely by evaluating the effect of each model on the quantity $(P_0^2 + P_{45}^2)$ determined for the estimation of the orientation parameters (see Eq. S12). Figure S2 shows the difference of values obtained in both models. It is found that this model can be reasonably used with typical differences of 10% found on $(P_0^2 + P_{45}^2)$ for $\delta$ ~ 100°.

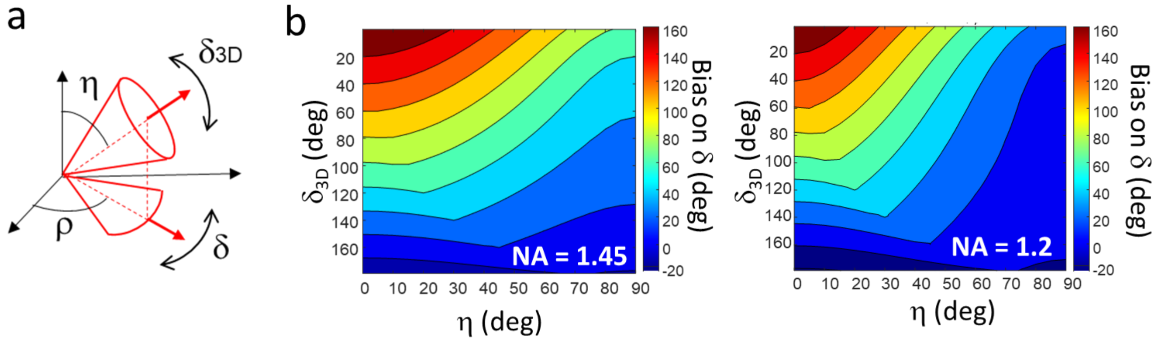


**Figure S1.** Retrieval bias on δ. (a) Schematic representation of a single molecule oriented in 3D by the mean orientation angles (ρ,η) and a wobbling cone angle of $\delta_{3D}$. The measured wobbling in 2D is δ. (b) Bias on δ (difference between the measured δ and the true value $\delta_{3D}$), under TIRF illumination and different detection NA conditions, as a function of the molecule tilt angle η (depicted in (a)).

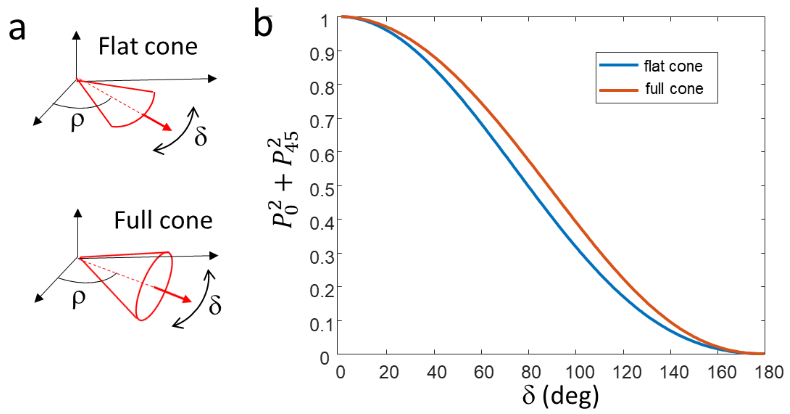


**Figure S2.** Cone model used for orientation parameter retrieval. (a) Approximated 2D 'flat cone' model used for the parameter retrieval, compared to a 'full cone' model. (b) The used parameter $P_0^2 + P_{45}^2$ for the retrieval of δ is depicted as a function of δ, in the case of a full cone model (red) and a flat cone model (blue). The difference between both models reaches 10% at δ = 100°.

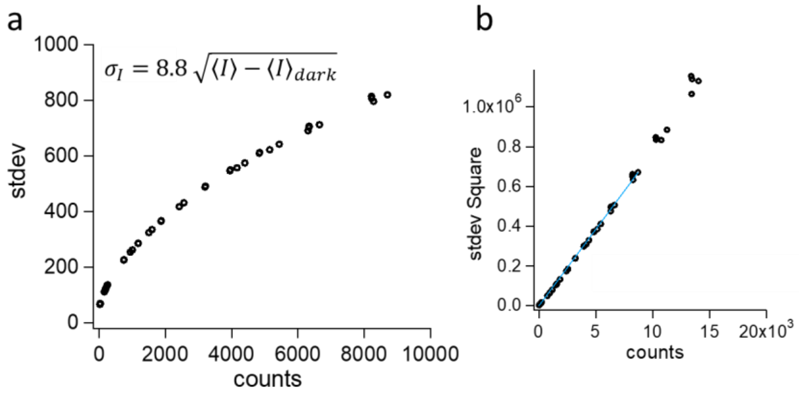


**Figure S3.** Camera noise estimation. (a) Dependence of the standard deviation $\sigma_I$ of the measured signal with respect to its mean value $\langle I\rangle$, measured on 500 samples (image regions of interest), using a camera gain of 300. The measurement was performed on homogeneous images produce by white light illumination of a piece of paper. Markers: experimental data. (b) Same data represented as $\sigma_I^2$. Blue line: linear fit, found to be of slope 77.5. The equation $\sigma_I = 8.8\sqrt{I - I_{dark}}$, is thus used as a noise model for the used camera in the present working conditions. An offset of $I_{dark} = 480$ counts is systematically removed to account for the camera electronic dark counts, irrespectively of the gain and exposure time. This offset is subtracted from all data acquisitions, including calibrations. The linearity of the camera response was moreover validated in the range of signals typically measured.

### Supplementary Information Note 2. Calibration factors in 4polar-STORM

Due to imperfections of the beam splitters as well as possible polarization leakages introduced by the optics of the detection path, correction factors need to be introduced in the estimation of the polarization factors $P_0$ and $P_{45}$. We denote $G_{BS}$ the factor accounting for the imperfect 50:50 reflection:transmission ratio of the first non-polarizing beam splitter. We also denote $G_0$ and $G_{45}$ the factors accounting for the unbalanced polarization split efficiency in the 0:90 and the 45:135 polarization channels respectively. $(G_{BS}, G_0, G_{45})$ are measured using a depolarized image (fluorescent solution) in which the ratios write:

$$G_{BS} = \frac{I_{45}+I_{135}}{I_0+I_{90}}\ ;\ G_0 = \frac{I_0}{I_{90}}\ ;\ G_{45} = \frac{I_{45}}{I_{135}} \qquad \text{Eq. S13}$$

We also account for possible polarization leakages between the 0:90 and 45:135 channels. The contributions are measured by polarizing the previous un-polarized image using a polarizer of known direction at the position of the back focal plane of the objective, before propagation through the dichroic filter and other propagation optics. This is performed using a controlled polarizer direction at the back focal objective plane (thin polarizer used, LPVISE100-A, Thorlabs). Denoting $G_1$ (resp. $G_2$) the leakage proportion factor of polarization channel 135° (resp. 45°) into the 45° channel (resp. 135°), and $G'_1$ (resp. $G'_2$) for the 0°/90° leakage, the final corrected expressions are:

$$P_0 = G_{BS} \cdot \frac{(1 + G'_1 - G'_2) \cdot I_0 - (1 - G'_1 + G_2) \cdot G_0 \cdot I_{90}}{(1 - G'_1 - G'_2) \cdot (I_0 + G_0 . I_{90})}$$

$$P_{45} = \frac{(1 + G_1 - G_2) \cdot I_{45} - (1 - G_1 + G_2) \cdot G_{45} \cdot I_{135}}{(1 - G_1 - G_2) \cdot (I_{45} + G_{45} . I_{135})}$$

Eq. S14

In an ideal optical system, $G_{BS} = G_0 = G_{45} = 1$ and $G_1 = G_2 = G'_1 = G'_2 = 0$. Deviations up to 0.2 from these numbers can be found in a real setup.

### Supplementary Information Note 3. Data processing algorithm of the 4polar-STORM method

The data processing is achieved by custom detection and analysis scripts written in Matlab, following a scheme summarized below. The scripts are based on previous work published in[10]. The previous polar-STORM algorithm, written for a two-image polarization split, has been adapted to a split into four images named (*img*$_0$*, img*$_{45}$*, img*$_{90}$*, Img*$_{135}$). Its goal is to estimate the polarization factors $P_0$ and $P_{45}$ for each detected single molecule, build a super-resolved image of these polarization factors, and reconstruct an orientation super-resolved image from them. The purpose of the 4polar-STORM algorithm is thus to retrieve, per molecule, its center point spread function (PSF) position coordinates in the 4 detected images ($i_0$, $i_{45}$, $i_{90}$, $i_{135}$), ($j_0$, $j_{45}$, $j_{90}$, $j_{135}$), its PSF radius ($r_0$, $r_{45}$, $r_{90}$, $r_{135}$), its localization precision ($\sigma_{loc,0}$, $\sigma_{loc,45}$, $\sigma_{loc,90}$, $\sigma_{loc,135}$) and its PSF amplitude ($\alpha_0$, $\alpha_{45}$, $\alpha_{90}$, $\alpha_{135}$), used to calculate the intensities ($I_0$, $I_{45}$, $I_{90}$, $I_{135}$) and thus the polarization factors $P_0$ and $P_{45}$ (see Eq. S7). Once these two factors are calculated, we deduce the orientation parameters for each single molecule (in-plane orientation angle ρ and wobbling angle value δ) using Eq. S12. This algorithm offers advantages as compared to a pure ratiometric calculation based on image registration, since the pairing is realized at the molecular level for each STORM stack recorded, without any requirement of pre-calibration experiment which may add additional positioning errors and therefore bias in the polarization factor estimation. This algorithm contains several sequential steps detailed below.

*Distortion correction.* A calibration sample (fluorescent nanobeads) is used to correct images from distortions. Fluorescent nanobeads of 100 nm in size (yellow-green Carboxylate-Modified FluoSpheres, ThermoFisher Scientific F8803) are immobilized on the surface of a poly-L-lysine coated coverslip and covered with a mounting medium (Fluoromount, Sigma F4680). A Fluorescent nanobeads image is used to estimate the spatial transformation (*tform*) to be used between the different polarization projections to correct any possible geometrical distortion, due mainly to the use of Wollaston prisms. We selected the quadrant $img_{90}$ as the reference image for the registration. The function *imregtform* (Matlab Imaging Processing toolbox) was used to retrieve the *tform* function for the other three quadrants ($img_0$, $img_{45}$ *and* $Img_{135}$). Then we correct these images using the transformation previously estimated. The spatial transformation was re-calculated only if the optical setup was modified. The correction was performed using the Matlab function *imwarp* under a linear interpolation (Matlab Imaging Processing toolbox).

*Detection and estimation*. As in the polar-STORM algorithm[10], single molecule localizations in image quadrants ($img_0$, $img_{45}$, $img_{90}$, $Img_{135}$) are based on a first detection step which uses a Generalized likelihood ratio test (GLRT) to identify the single molecule candidates for the STORM image reconstruction, as detailed in[12]. This detection step uses a given fixed Gaussian shape for the theoretical PSF (starting from an initial guess radius of $r$ = 1.3 pixels), a spatial sliding detection window ($w_s$) and a limit probability of false alarm (PFA)[12], which defines a threshold limit (calculated empirically based on Monte Carlo simulations) above which any signal can be statistically considered as having a different origin than noise. A value PFA ≤ $10^{-6}$ is set to guarantee a probability of false alarm (PFA) of less than 1 pixel per image, which ensures a fraction of the detected single molecules close to 100% for a signal to noise ratio (SNR) higher than 20dB[12]. After the candidates have been detected by the GLRT algorithm, the amplitude, radius and position of their Gaussian PSF are estimated on all four quadrants at the subpixel level, based on a Maximum likelihood (ML) estimation using a Gauss-Newton regression. This regression uses the GRLT obtained values of radius and position as initial parameters. The localization accuracy ($\sigma_{loc}$) is estimated from a computation of the Cramer-Rao bound (CRB) limit[12], and given for all quadrant images.

*Estimation of the translation vector between image quadrant pairs: ($img_0$ - $img_{90}$), ($img_{45}$ - $Img_{135}$) and ($img_0$ - $img_{45}$).* To estimate the translation vector between the different images and associate each detected PSF to a given molecule, it is possible to use the registration from the bead sample used for the distortion-correction step above. This type of registration is however limited by the camera pixel size, image quality and stability of the optical system. It also implies the use of interpolation methods during the image subsampling. The 4polar-STORM software rather directly calculates the translation vector using the detected molecules themselves, which are localized with high precision. The distance of each molecules' images in the quadrant pairs *($img_0$ - $img_{90}$), ($img_{45}$ - $Img_{135}$)*, and *($img_0$ - $img_{45}$)* *c*reated by the Wollaston polarization beam splitter prisms is represented by three vectors ($\vec{u}_{0-90}$ , $\vec{u}_{45-135}$, $\vec{u}_{0-45}$). The knowledge of these translational vectors is required for the detection of the molecule-pairs present in a STORM image stack. To estimate the vectors, we perform a statistical estimation by using the 100-1,000 first frames of the STORM recorded stack[10]. First, all possible vectors joining two molecules of the images are calculated and the squared norms of the differences between all the obtained vectors are calculated. This leads to a statistical distribution, within which only the candidates whose difference is below 4.2 times the obtained standard deviation are kept (pure significant test assuming that the difference between norms follows a Gaussian distribution, which guarantees a 95% confidence level to have similar directions between the selected pairs of PSF images). Second, an additional selection is performed to determine the most optimal vector. This second step compares the obtained vector squared-norms and keeps the largest ensemble of similar squared-norms in this population. For this a sub-optimal detection is run (the only hypothesis being that the error on position is Gaussian), keeping errors only below a threshold that guaranties a 95% confidence level within the obtained distribution.

*Association of molecule pairs.* After the three optimal vectors ($\vec{u}_{0-90}$, $\vec{u}_{45-135}$, $\vec{u}_{0-45}$) are calculated, pairs of molecules along these directions are coupled by selecting the nearest neighbor to the expected position at a vector distance from the reference quadrant, within a distance tolerance corresponding to the localization precision. First, molecules from the images pairs ($img_0$ - $img_{90}$) and ($img_{45}$ - $img_{135}$) are coupled separately over the full STORM stack (typically 30 000 – 50 000 images), then the identified couples are associated using $\vec{u}_{0-45}$.

*Estimation of molecular parameters*. The pairing of all molecules allows the reconstruction of a polarization STORM image based on the parameter position ($i_0$, $i_{45}$, $i_{90}$, $i_{135}$), ($j_0$, $j_{45}$, $j_{90}$, $j_{135}$), PSF radius ($r_0$, $r_{45}$, $r_{90}$, $r_{135}$), localization precision ($\sigma_{loc,0}$, $\sigma_{loc,45}$, $\sigma_{loc,90}$, $\sigma_{loc,135}$) and PSF amplitude ($\alpha_0$, $\alpha_{45}$, $\alpha_{90}$, $\alpha_{135}$). Only molecules that were presented in all four-quadrants were considered for analysis to avoid bias in the orientation and intensity estimation.

*Calculation of polarization factors $P_0$ and $P_{45}$.* Polarization factors $P_0$ and $P_{45}$ (see Supplementary Note 1) are calculated based on the integrated intensities $I_0$, $I_{45}$, $I_{90}$ and $I_{135}$, which are calculated from ($\alpha_0$, $r_0$), ($\alpha_{45}$, $r_{45}$), ($\alpha_{90}$, $r_{90}$) and ($\alpha_{135}$, $r_{135}$):

$$I = 2\sqrt{\pi} r \alpha \quad \text{Eq. S15}$$

After intensities are estimated, the polarization factors ($P_0, P_{45}$) are deduced, accounting for the calibration correction factors as detailed in Eq. S14 (see Supplementary Note 2).

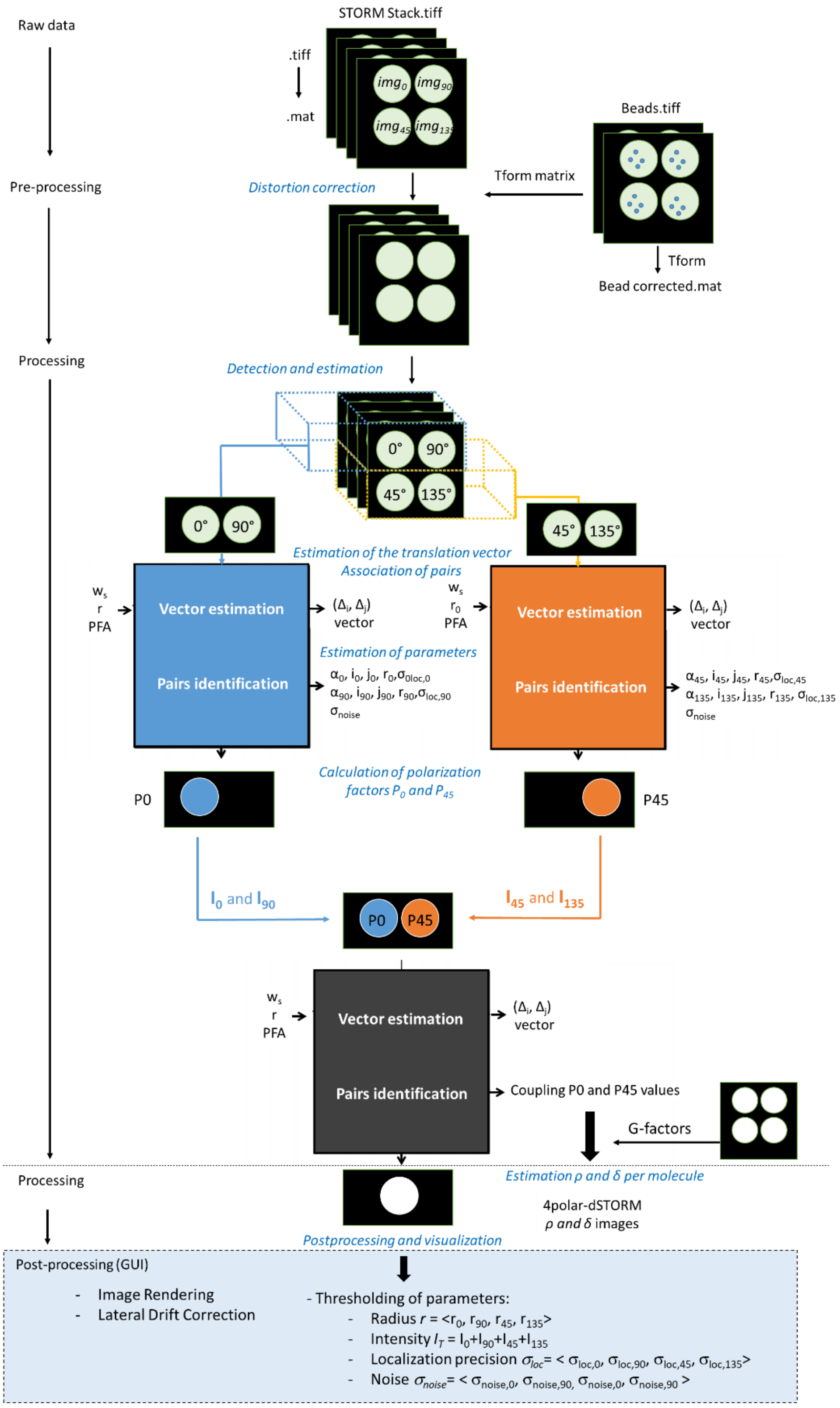


**4polar-STORM algorithm flowchart.** In the pre-processing step, the raw data (.tiff image stack files) is transformed into ".mat" files to be further spatially corrected by the bead calibration in Matlab. The distortion-corrected stack files are processed for each Wollaston polarized arm (in blue, the pair 0°-90°, and in orange, the pair 45°-135°). On each arm, the 4polar-STORM algorithm is applied: GLRT is applied to find the best translation vector, which is further used to identify coupled blinking pair events. A multi-parametric Gaussian-Newton fit is performed to retrieve quantitative parameters of the blinking particles, such as position (i,j), localization precision ($\sigma_{loc}$), the amplitude (α), radius (r), and noise ($\sigma_{noise}$). A vector estimation and pair identification (coupling) is done to group the information into polarization factor ratios P0 and P45. The last step of the data treatment is to calculate the orientation parameters for each blinking particle in the reconstructed image. For that, the G-factors (i.e., the intensity calibration correction) are applied. Post-processing is performed using a derivation of PALMSiever[13]. The most common post-processing steps used for 4polar-STORM are: (1) lateral drift correction (based on cross-correlation with the localization themselves), (2) choice of image rendering, (3) parameter thresholding for robust orientation parameter estimation, and (4) choice of the stick representation.

*Estimation of ρ and δ per molecule*. After the polarization factors are calculated, *ρ* and *δ* are estimated based on the expressions given in Eq. S12. To solve the determination of *δ,* an interpolation is performed using the function "*interp1*" of Matlab.

*Postprocessing and visualization.* Postprocessing is performed in a modified version of PALMsiever[13], a visualization and analysis platform for single-molecule localization microscopy implemented in Matlab. It uses the software package DIPimage (https://diplib.org/DIPimage). PALMsiever includes a plugin for drift correction using cross-correlation. For this work, we used two rendering modalities: histogram + Gaussian filter and Kernel Density Estimation (KDE)[14]. KDE is a smoothing version of histogram, with the smoothing kernel bandwidth estimated from the molecule density. Our version of PALMsiever of the 4polar-STORM modality includes the estimation of *ρ* and *δ*, as well as their graphical representation as sticks. In this representation, *ρ* is depicted as the orientation angle of sticks (with respect to the horizontal axis of the image), and δ or *ρ* are depicted as the colors of the sticks. The sticks are displayed over a black-and-white image representing the super-resolved STORM image, which uses the localization of all detected molecules. 4polar-STORM allows to use the default parameter filters of PALMsiever for the image representation, filtering for instance molecule populations by their density, intensity and localization precision. An example of 4polar-STORM representation is shown below.

*Representation of ρ and δ in STORM images*. The final 4polar-STORM representation consists in depicting, over a STORM image background, one stick per single molecule detected, whose orientation relative to the horizontal axis is *ρ,* and whose color is either encoding *ρ* or *δ*. In the chosen representation, we plot sticks with largest *δ* values above sticks with lowest ones, in order to better visualize the presence of highly wobbling populations in red.

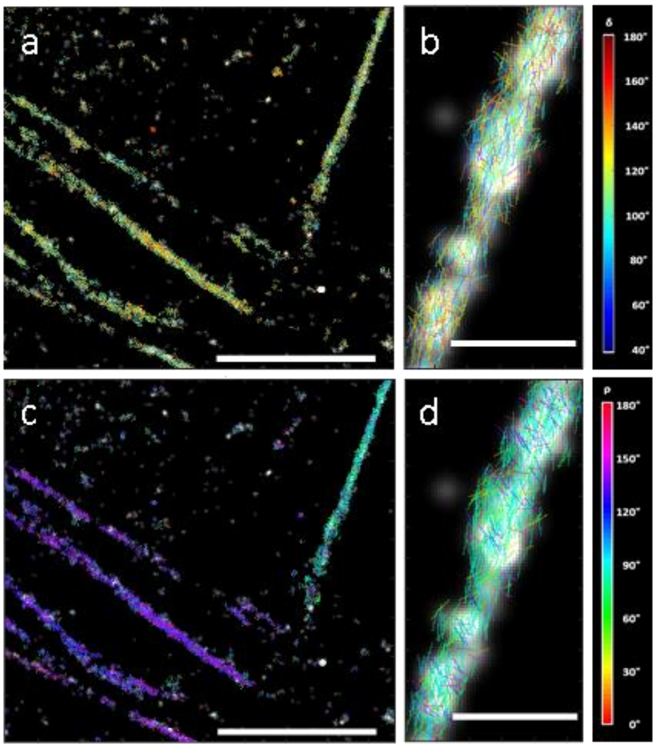


**4polar-STORM stick representations of (δ,ρ).** Example of super-resolved image of δ (a,b) and ρ (c,d) using 4polar-STORM on stress fibers of a U2OS cell stained for F-actin (AF488-phalloidin). Stick orientation is based on ρ. Colors of the sticks correspond to δ (a,b) and ρ (c,d). Scale bars: 5 µm (a and c) and 500 nm (b and d). Gaussian blurring size: 39 nm. A density filter was applied to remove isolated spots. Rendering pixel size: 12.42 nm (b,d), 24.86 nm (a, c).

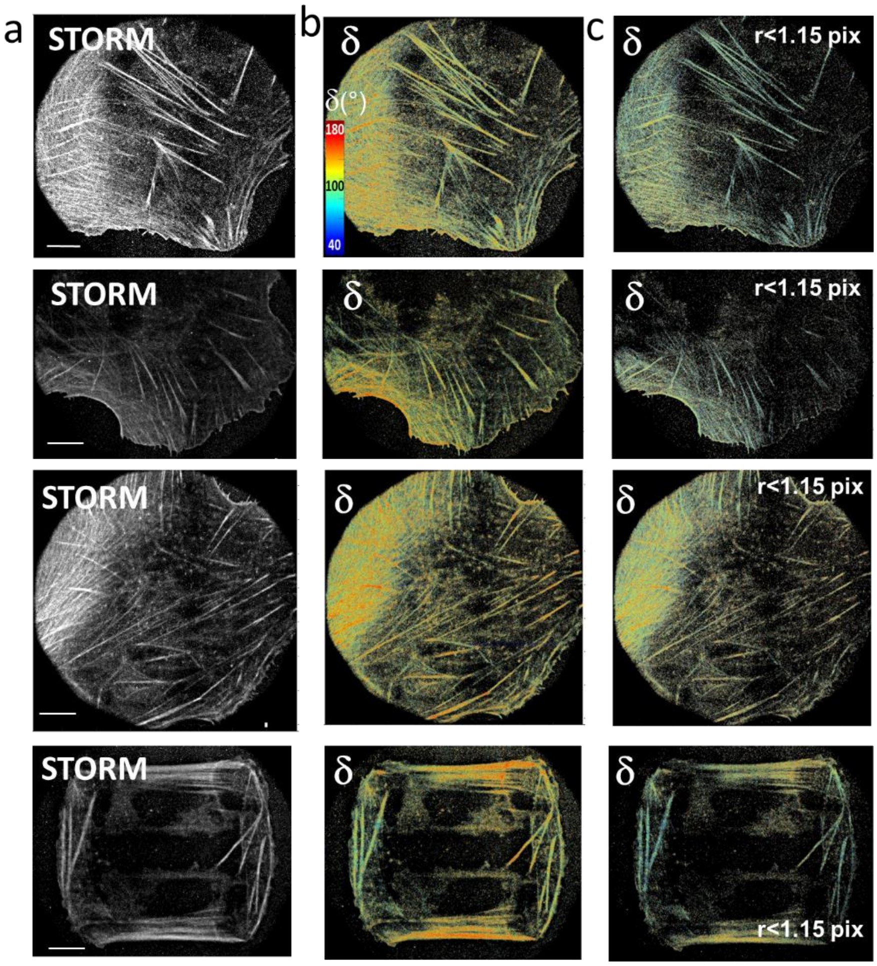


**Figure S4.** 4polar-STORM $\delta$ images of F-actin in fixed U2OS cells labelled with AF488-phalloidin. (a) STORM images. (b) Corresponding $\delta$ images with no thresholding of the detection parameters. (c) Corresponding $\delta$ images keeping only molecules for which the PSF radius r is below 1.15 pixels (~150 nm). $\delta$ color scales are the same for all images. Scale bars 6.5 $\mu$m.

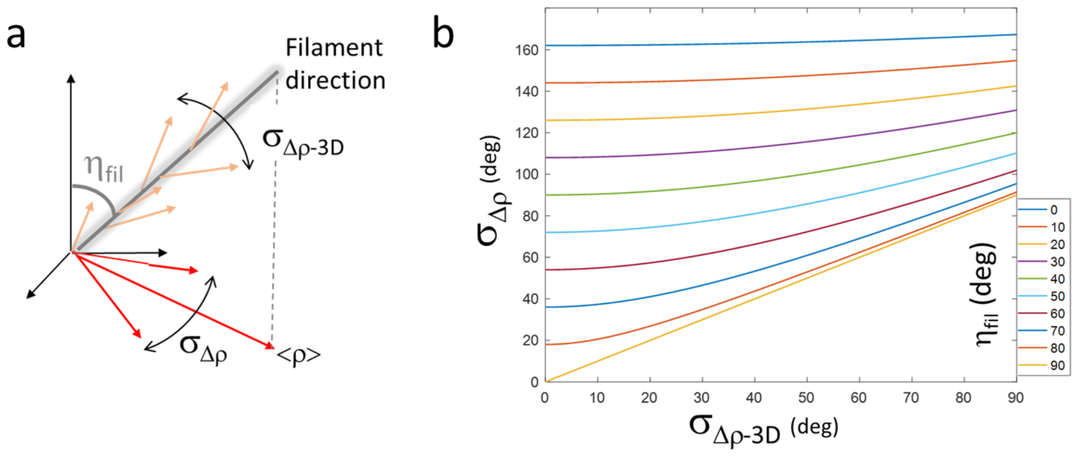


**Figure S5.** Retrieval bias on $\sigma_{\Delta\rho}$. (a) Schematic representation of an actin filament tilted off-plane by an angle $\eta_{fil}$, with a distribution of single molecule orientations represented by $\sigma_{\Delta\rho\text{-3D}}$. The measured distribution of projected orientations in 2D is represented by $\sigma_{\Delta\rho}$. (b) The graph shows the measured $\sigma_{\Delta\rho}$ as a function of the true value $\sigma_{\Delta\rho\text{-3D}}$, for different filament off-plane angles $\eta_{fil}$. This calculation is derived from purely geometrical considerations, considering cone apertures projected in the sample plane after a rotation is applied to them to simulate off-plane tilt.

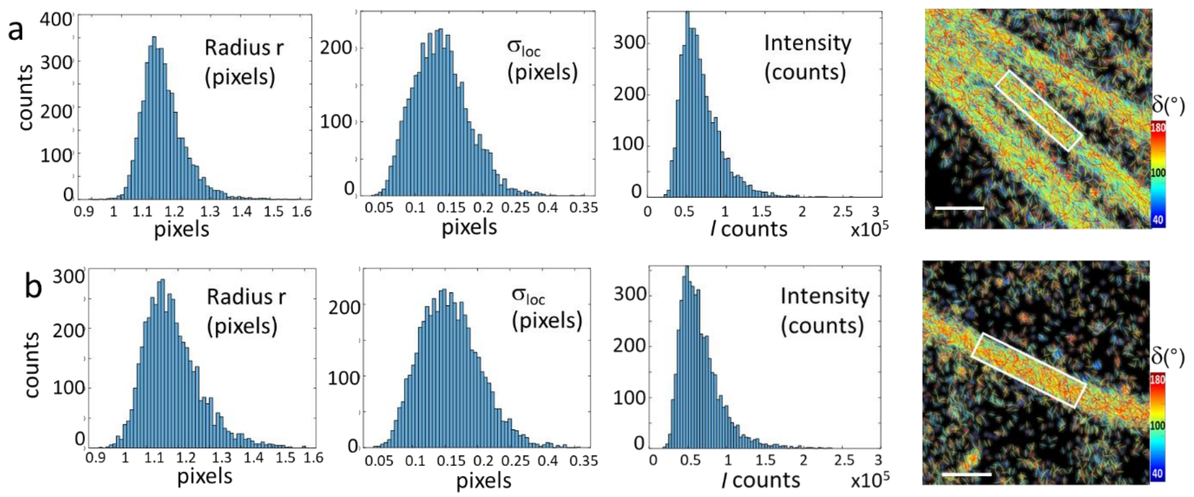


**Figure S6.** Statistics on detection parameters in 4polar-STORM imaging of F-actin in stress fibers in cells. Histograms of the detection parameters in different types of stress fibers in U2OS cells labelled with AF488-phalloidin (shown as $\delta$ stick images). The histograms depict values measured for all molecules present in the regions of interest shown as white rectangles. (a) In-plane ventral SF region. (b) Focal adhesion region, for which the distributions of the radius r and localization precision $\sigma_{loc}$ become wider. Scale bars: 800 nm.

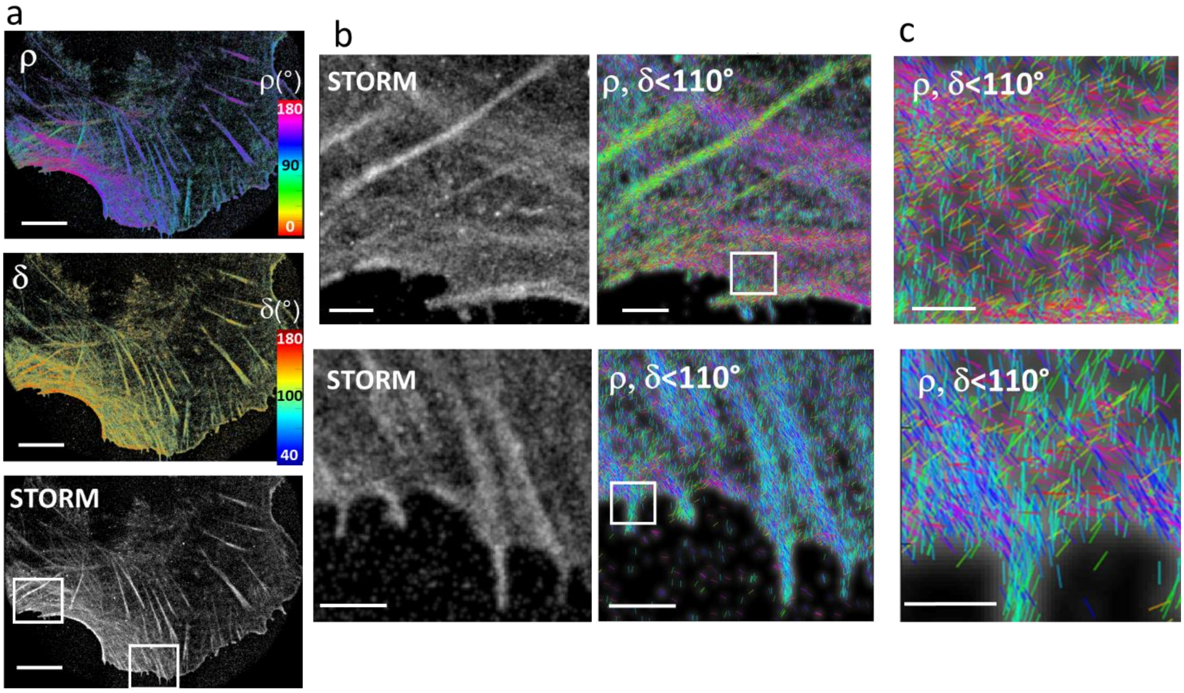


**Figure S7.** 4polar-STORM imaging of F-actin in cells, selecting in-plane actin filament populations. (a) Large field of view images of ρ and δ sticks as well as the corresponding single molecule localization STORM image of a U2OS cell labelled with AF488-phalloidin. (b) zoomed regions (see squares in (a)) depicting STORM and ρ-stick images for in-plane molecules only (δ < 110°). (c) stronger zoom (see squares in (b)) depicting ρ sticks for in-plane molecules only (δ < 110°). Scale bars (a) 6.5 μm; (b) 1.3 μm; (c) 500 nm.

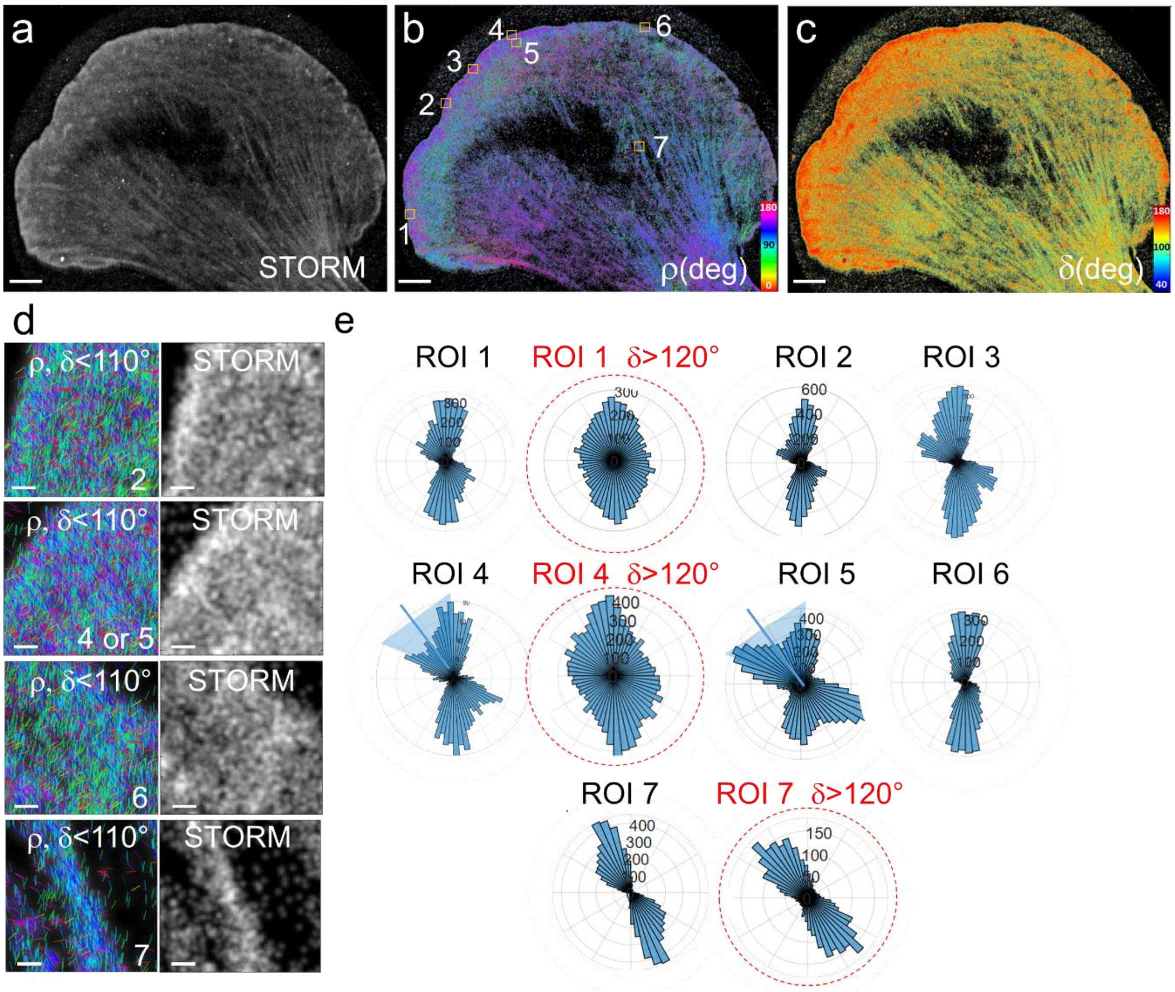


**Figure S8.** 4polar-STORM imaging of actin filament organization in lamellipodia. (a) Single molecule localization STORM image of a B16 cell labelled with AF488-phalloidin. (b) Corresponding 4polar-STORM ρ stick image with color-coded orientation measurements. (c) 4polar-STORM δ stick image with color-coded wobbling angle measurements. (d) Examples of ρ stick images showing molecules with $\delta < 110°$ and corresponding STORM images in selected ROIs (squares in (b)). ROIs 1-6, regions in the lamellipodium; ROI 7, SF. (e) Polar-plot histograms of ρ for the regions shown in (b). The condition $\delta < 110°$ is used, except for red-circled histograms for which $\delta > 120°$ molecules are selected. Scale bars (a-c), 4 μm; (d), 260 nm.

### Supplementary Information References

6. C. R. Cantor and P. R. Schimmel. *Biophysical Chemistry. Part II: Techniques for the Study of Biological Structure and Function*.
